# HER2 overexpression initiates breast tumorigenesis non-cell-autonomously by inducing oxidative stress in the tissue microenvironment

**DOI:** 10.1101/2023.08.25.554770

**Authors:** Sevim B. Gurler, Oliver Wagstaff, Lili Dimitrova, Fuhui Chen, Robert Pedley, William Weston, Ian J. Donaldson, Brian A. Telfer, David Novo, Kyriaki Pavlou, George Taylor, Yaqing Ou, Kaye J. Williams, Andrew Gilmore, Keith Brennan, Ahmet Ucar

## Abstract

HER2 is considered as a driver oncogene responsible for the HER2+ subtype of breast cancer. However, it is still unclear how HER2 induces the oncogenic transformation of breast cancer stem cells (BCSCs) and initiates tumorigenesis during premalignant stage breast cancer. Here, we used clinical samples and mouse models of HER2+ breast cancer to demonstrate that neither BCSCs nor their cell-of-origin express HER2/Neu in early-stage breast tumors. Instead, our results demonstrate that Neu overexpression results in the transformation of BCSCs in a non-cell-autonomous manner via triggering DNA damage and somatic mutagenesis in their Neu-negative cell-of-origin. This is caused by the increased oxidative stress in the tissue microenvironment generated by altered energy metabolism and increased reactive oxygen species levels in Neu-overexpressing mammary ducts. Therefore, our findings illustrate a previously unrecognized mechanism of HER2-induced breast tumor initiation *in vivo* with potential impacts on future preventive treatments for HER2+ premalignant breast cancer.

## INTRODUCTION

Breast cancer is the most common cancer and one of the leading causes of cancer-related deaths worldwide^1^. Around 15–20% of malignant breast tumors are defined as HER2+ subtype and characterized by the overexpression and/or amplification of the *HER2/Erbb2* oncogene. Although the mechanisms of HER2 signaling in breast tumor cells have been extensively studied, our current knowledge on HER2-driven tumorigenesis is still incomplete, especially for its role in pre-malignant stages of breast tumorigenesis.

Clinical observations suggest an enigmatic role for HER2 during the malignant transition from Stage-0 ductal carcinoma *in situ* (DCIS) to invasive ductal breast carcinoma^2, 3^. HER2 overexpression/amplification is rarely detected in premalignant lesions of atypical ductal hyperplasia, whereas up to 70% of high-grade DCIS are characterized as HER2+. The dramatic difference between the prevalence of HER2+ DCIS versus HER2+ invasive breast cancer raises clinically important questions on the functional significance of HER2 in premalignant breast cancer.

Breast cancer stem cells (BCSCs) reside at the apex of tumor cell ontogeny due to their stem cell-like stemness features and have the unique tumor-initiating abilities. Therefore, they are presumed to be the key drivers of tumor progression, metastasis, treatment resistance, and tumor recurrence^4, 5^. It is likely that tumor evolution is mechanistically driven by the accumulation of somatic mutations in BCSCs or their cell-of-origin. However, we currently have a very limited understanding of how BCSCs arise during tumor initiation and further evolve during tumor progression. Furthermore, it is still under debate whether BCSCs are a distinct tumor cell type or a tumor cell state within the tumor cell ontogeny, as it was shown that cytoablative treatments can induce dedifferentiation of non-stem tumor cells into BCSCs^6^. Notwithstanding these controversies, little is known about how an oncogene expression, such as HER2, can result in the transformation of BCSCs from their cell-of-origin during premalignant stages of breast cancer.

Here, we used clinical DCIS samples and two distinct mouse models of HER2+ breast cancer to show that BCSCs do not express HER2 during early stages of tumorigenesis. Furthermore, the lineage tracing analysis revealed that BCSCs arise from cells that never expressed HER2/Neu, thus suggesting a non-cell-autonomous effect of HER2/Neu overexpression on the cell-of-origin of BCSCs. Histological analyses of MMTV-Neu mammary glands showed ectatic duct formation prior to tumor development and a ductal lineage dependency of early-stage tumorigenesis. Proteomics and metabolomics analyses of MMTV-Neu mammary ducts prior to the formation of histologically apparent tumors demonstrated an altered activity in energy metabolism, which consequently results in increased oxidative stress in the tissue microenvironment that leads to DNA damage and mutagenesis not only in the HER2/Neu-expressing cells, but also in the HER2/Neu-negative cells. Thus, our results suggest that BCSCs may transform from their HER2/Neu-negative cell-of-origin as a consequence of the mutagenic oxidative microenvironment caused by the altered metabolism of HER2/Neu-overexpressing mammary epithelial cells during HER2-induced breast tumorigenesis.

## RESULTS

### Neither BCSCs nor their cell-of-origin express HER2/Neu oncogene during early stages of HER2/Neu-driven breast tumorigenesis

Earlier studies using human breast cancer cell lines demonstrated that a subpopulation of BCSCs may express HER2^7–10^. However, there are no conclusive studies available that determined the HER2 expression status of BCSCs within DCIS. Thus, we analyzed clinical samples of DCIS to characterize their BCSCs. Flow cytometry analyses of the lineage (CD31, CD45, CD235a)-negative DCIS cells for EpCAM versus HER2 expression revealed a considerable level of inter-patient heterogeneity (Supplementary Figure 1). We categorized these 15 DCIS samples into 3 putative groups based on the frequency of HER2^pos^ cells (Supplementary Figure 1). Previous studies on normal human breast samples demonstrated that luminal epithelial lineage forms the EpCAM^high^ population, whereas mammary epithelial stem cells (MaSCs) reside within the EpCAM^neg/low^ population^11, 12^. Thus, we sorted three subpopulations of cells from the HER2-enriched and HER2-low DCIS samples and plated them in mammosphere culture to determine which populations might contain BCSCs (Figure 1A– B). For all DCIS samples analyzed, EpCAM^high^ cell populations formed only acini structures, resembling to those formed by luminal progenitors isolated from normal mammary glands^13,14^. In contrast, mammospheres were formed solely by the EpCAM^neg/low^ cell populations and they were most frequently observed within the cultures of the HER2^neg^ subpopulation (Figure 1C). These results demonstrate that BCSCs reside either predominantly or exclusively within the EpCAM^neg/low^;HER2^neg^ subpopulation of cells in DCIS.

**Figure 1.**
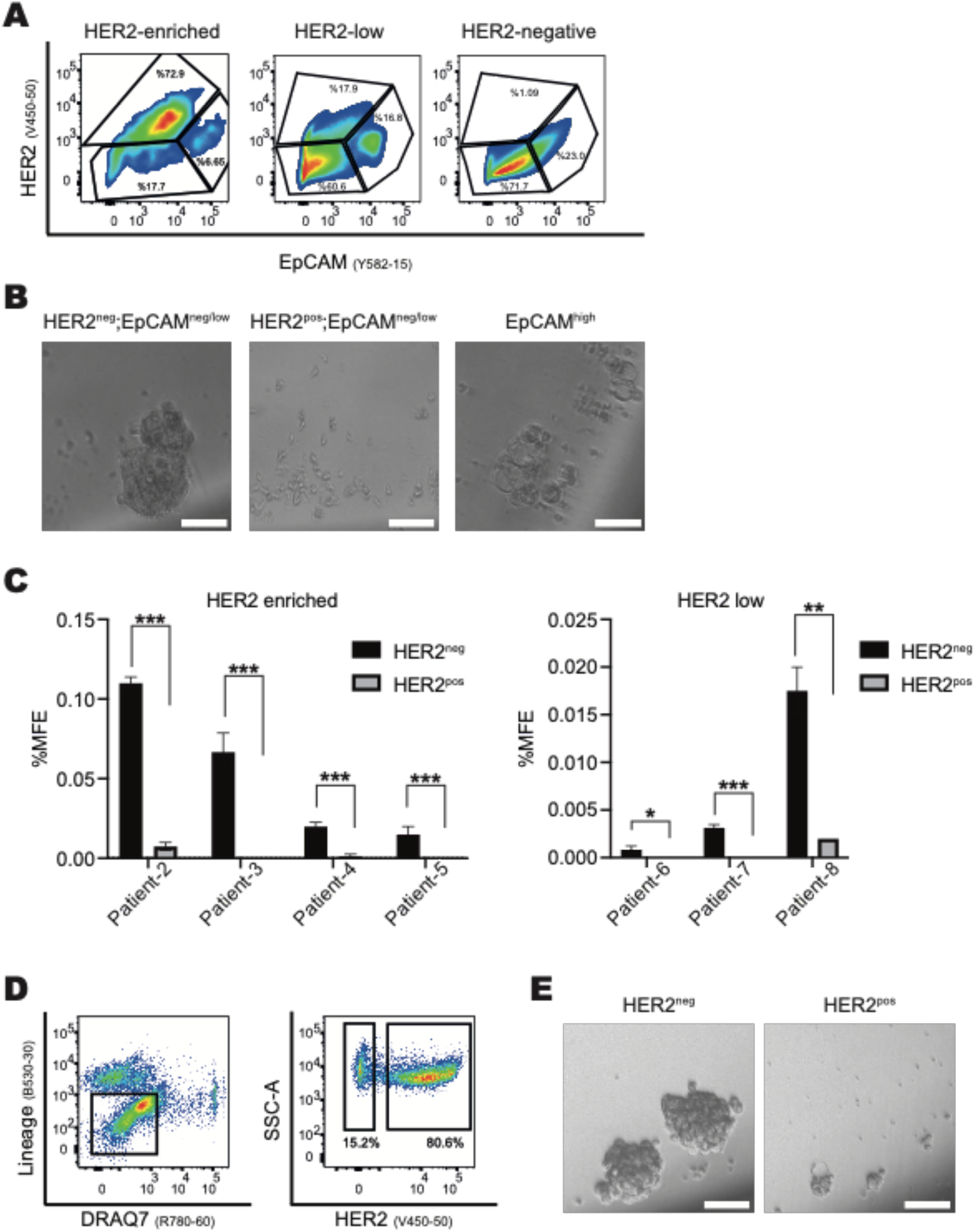
BCSCs do not express HER2 in HER2+ DCIS or early-stage HER2+ murine tumors. **A)** Flow cytometry analysis identifies three putative subgroups of DCIS based on the abundance of their HER2^pos^ cells. Shown here are pseudo-colored dot plots representing HER2-enriched, HER2-low and HER2-negative DCIS subgroups for the HER2 versus EpCAM expression within their lineage-negative cell populations (see Supplementary Figure 1). Depicted gates represent three subpopulations of DCIS cells that were sorted for analysis. **B)** Mammosphere culture of DCIS subpopulations sorted using the gates shown in (**A**). Micrographs shown are representative of all analyzed DCIS samples and demonstrate that (Lin^neg^;EpCAM^high^) cells form small acini structures, (Lin^neg^;HER2^neg^;EpCAM^neg/low^) form mammospheres, and (Lin^neg^;HER2^pos^;EpCAM^neg/low^) cells do not form any structures. Scale bars represent 50 um. **C)** Quantification of the mammosphere-forming efficiency (%MFE) for the Lin^neg^;HER2^neg^;EpCAM^neg/low^ and Lin^neg^;HER2^pos^;EpCAM^neg/low^ subpopulations of DCIS cells shows that BCSCs are predominantly or exclusively present in Lin^neg^;HER2^neg^;EpCAM^neg/low^ subpopulation. Values represent mean ± SEM of 2 to 8 technical replicates. *:p<0.05; **:p<0.01; ***:p<0.0001; by multiple unpaired t-test comparisons. **D)** Representative dot plots for the flow cytometry analysis of MMTV-rTTa;TetON-HER2 tumors (n=3). Single cells that are negative for lineage markers (CD31, CD45, TER119) and dead cell marker (DRAQ7) are gated (left plot) for visualizing their HER2 expression levels (right plot). **E)** Mammosphere culture of Lin^neg^;HER2^neg^ and Lin^neg^;HER2^pos^ subpopulations of MMTV-rTTa;TetON-HER2 tumor cells sorted as shown in (**D**) demonstrates that BCSCs reside exclusively in the Lin^neg^;HER2^neg^ subpopulation. Scale bars represent 50 um.

To determine whether a similar observation can be made in a mouse model of HER2+ breast cancer, we employed the MMTV-rTTa;TetON-HER2 double-transgenic mouse line^15^ that enables doxycycline-inducible *HER2/ERBB2* overexpression in murine mammary glands. We induced HER2 overexpression in 15-weeks old double transgenic females that consequently developed palpable mammary tumors within 4 to 5 months of continuous doxycycline treatment. These tumors were then analyzed for their HER2 expression by flow cytometry. Although most lineage-negative tumor cells were identified as HER2^pos^, there was a population of 15% to 30% of tumor cells that were HER2^neg^ (Figure 1D). When sorted and plated in mammosphere culture, only the HER2^neg^ cell subpopulation have formed mammospheres (Figure 1E), indicating that BCSCs reside exclusively within the HER2^neg^ cell subpopulation in this mouse model, similar to our observations in clinical DCIS samples.

We then asked whether BCSCs may arise from a HER2-expressing cell-of-origin, despite not expressing HER2 themselves. To address this question, we employed a lineage tracing approach using another mouse model of HER2+ breast cancer, MMTV-Neu-IRES-Cre (MMTV-NIC)^16^. We crossed this line with the CRE-recombinase dual reporter R26R-mT/mG mouse line^17^ to analyze tumors formed by MMTV-NIC;R26R-mT/mG double transgenic females. In these mice, tdTomato expression marks the Neu^neg^ lineage, which consists of cells that neither express the Neu oncogene themselves nor are derived from a Neu-expressing cell. In contrast, GFP-expressing cells constitute the Neu^pos^ lineage, which are those that express the Neu oncogene themselves and/or are derived from a Neu-expressing cell-of-origin. As expected, most cells within the tumors of these double-transgenic mice were GFP^+^, whereas tdTOM^+^ cells constituted only a small subpopulation of 0.4% to 6% of tumor cells (Figure 2A). We then sorted and plated these cells in mammosphere culture. Interestingly, mammospheres were only formed by tdTOM^+^ tumor cells (Figure 2B), suggesting that BCSCs reside exclusively within the Neu^neg^ lineage. Next, we analyzed the mammary glands of 12-weeks old MMTV-NIC;R26R-mT/mG mice that had no apparent tumors in their glands to determine whether the MMTV promoter is active in MaSCs in this mouse line. Our results demonstrated that mammosphere-forming MaSCs also resided within the tdTOM^+^/Neu^neg^ lineage (Figure 2A,B), suggesting that MMTV promoter does not drive the Neu expression in MaSCs in this mouse model.

**Figure 2.**
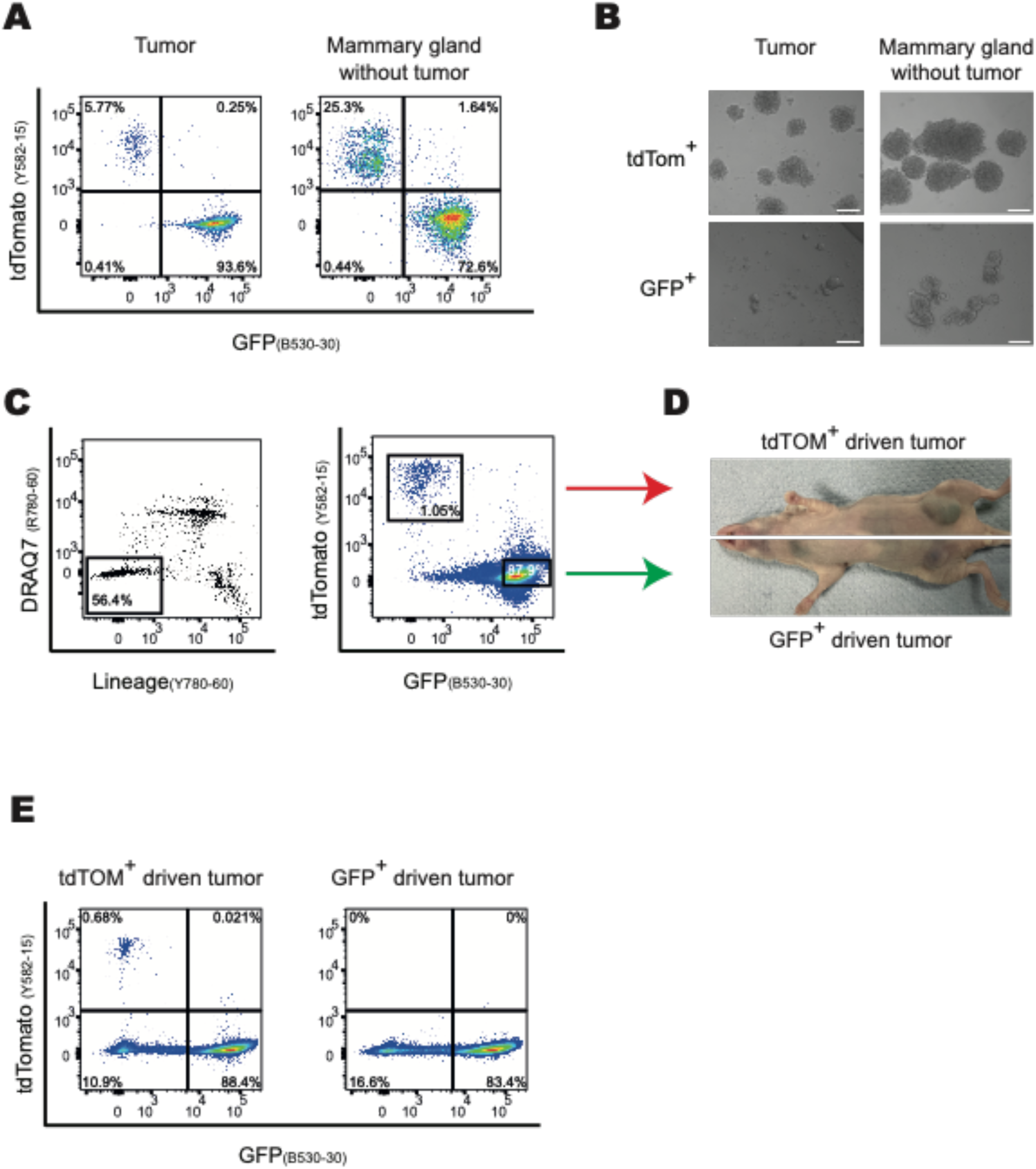
BCSCs reside in the Neu^neg^ cell lineage in MMTV-Neu tumors. **A)** Flow cytometry analyses of MMTV-NIC/+;R26R-mT/mG tumors (n=10) and mammary glands (n=3) reveal that Neu^pos^ and Neu^neg^ lineage cells can be distinguished as GFP^+^ and tdTOM^+^ cells, respectively. **B)** Mammosphere cultures of GFP^+^ and tdTOM^+^ cells sorted from MMTV-NIC/+;R26RmT/ mG tumors and mammary glands as shown in (A) demonstrate that mammosphere-forming cells reside exclusively in the Neu^neg^ lineage. Scale bars represent 50 um. **C)** Representative dot plots used for sorting tdTOM^+^ and GFP^+^ cells from MMTVNIC/+; R26R-mT/mG tumors for transplantation. Single cells that are negative for lineage markers (CD31, CD45, TER119) and dead cell marker (DRAQ7) are gated (left plot) for identifying tdTOM^+^ and GFP^+^ tumor cells (right plot). **D)** Pictures of the transplanted nude mice developing tumors in their No:4 inguinal mammary glands upon orthotopic transplantation of 50,000 tdTOM^+^ or GFP+ tumor cells that were sorted as shown in (C). **E)** Flow cytometry analysis of tumor transplants for the tdTomato versus GFP expression in their Lin^neg^ population reveals that tdTOM^+^ cells can generate GFP^+^ cells during tumor development. Note that double-negative cells are the host cells originating from nude mice. (n=2 transplant for each tumor type).

Next, we investigated whether Neu^neg^ lineage cells can form tumors upon transplantation, a characteristic feature of BCSCs. GFP^+^ and tdTOM^+^ cell populations were sorted from MMTV-NIC;R26R-mT/mG tumors and orthotopically transplanted into the mammary fat pads of immune-deficient nude mice (Figure 2C). We observed that both cell fractions formed tumors upon transplantation (Figure 2D). Importantly, flow cytometry analyses of these tumors revealed that tdTOM^+^/Neu^neg^ lineage cells were able to generate GFP^+^ cells that constituted most of the tumor cell population, while still maintaining a small subpopulation of tdTOM+ cells (Figure 2E). This suggests that BCSCs reside in the Neu^neg^ lineage and can generate Neu^pos^ lineage cells during tumor formation.

Taken together, our results indicate that neither BCSCs nor their cell-of-origin express the HER2/Neu oncogene during early-stages of HER2/Neu-induced breast tumorigenesis. This may imply that the transformation of BCSCs from their cell-of-origin must rely on a non-cell-autonomous effect of HER2/Neu-overexpression in mammary epithelia.

### Neu-driven tumorigenesis displays ductal lineage dependency *in vivo*

In MMTV-NIC mouse line, all nulliparous females develop mammary tumors at around 4–5 months of age^16, 18^. Although all five pairs of their mammary glands are prone to develop tumors, palpable tumors often develop quicker in their thoracic and cervical glands compared with their inguinal glands. Thus, to visualize the earlier stages of mammary tumorigenesis, we performed whole-mount staining on No:4 inguinal glands of MMTV-NIC females that had palpable tumors only in their thoracic or cervical glands. In all No:4 glands analyzed, we observed multi-nodal early-stage tumor formation being restricted to only a few primary ducts (Figure 3A), and these cancerous growths were observed to be protruding from mammary ducts rather than being located within the ducts (Figure 3B). To confirm whether the whole mammary ductal tree, rather than only the tumor-bearing primary ducts, are developed from Neu-expressing cells in this mouse model, we analyzed the No:4 inguinal glands of 4 to 5 months old MMTV-NIC;R26R-mT/mG double transgenic mice. A similar beads-on-string pattern of early-stage GFP^+^ tumors were observed only in few ducts in these glands, although the entire ductal tree was GFP^+^ (Figure 3C).

**Figure 3.**
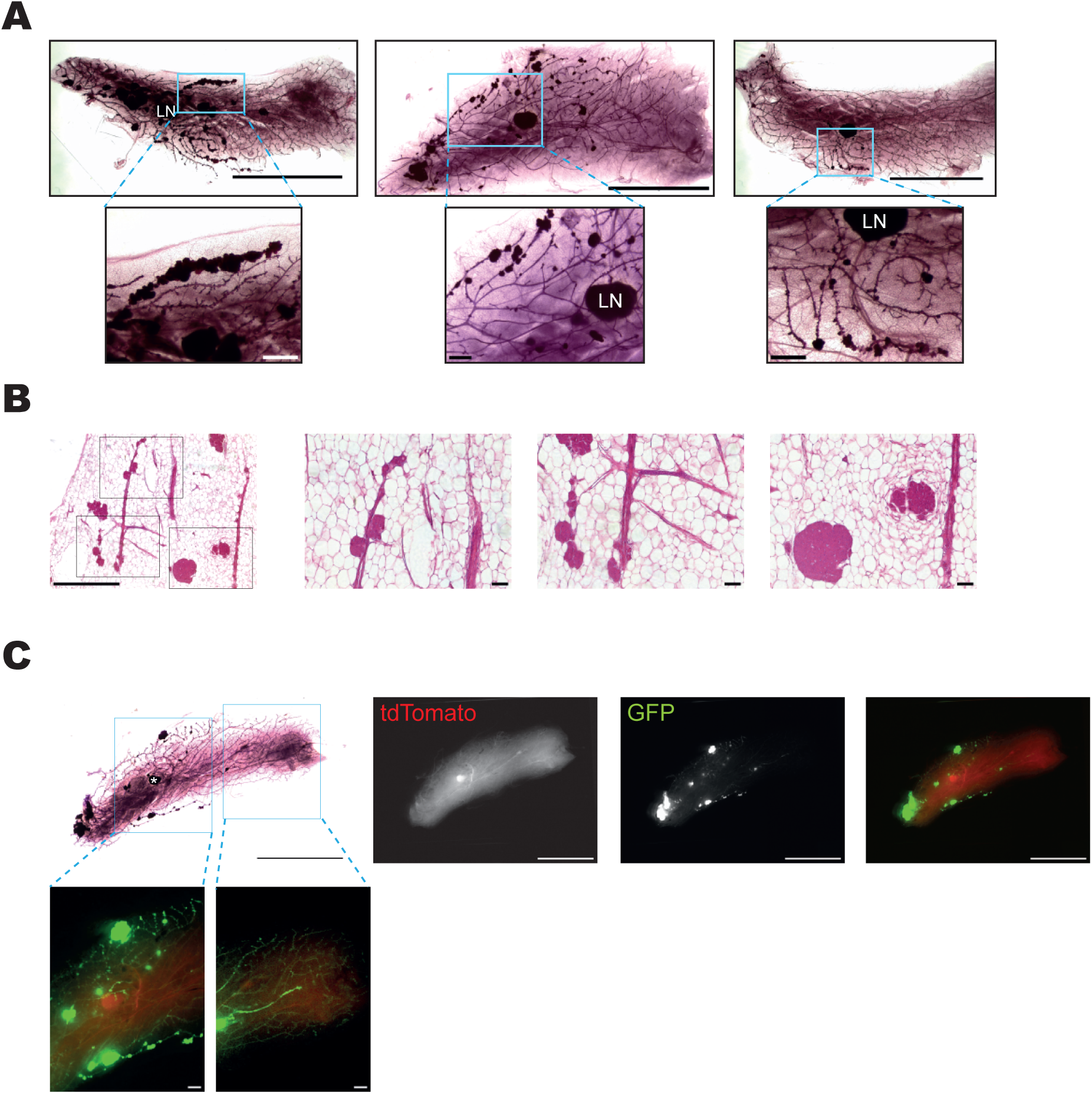
Neu-driven early-stage mammary tumorigenesis displays ductal-lineage dependency. **A)** Whole-mount staining results of the No:4 inguinal glands of three adult MMTV-NIC mice demonstrates a beads-on-string pattern of early-stage mammary tumourigenesis indicating a ductal lineage–dependency of tumor development. LN: lymph node. Scale bars represent 1 cm and 1 mm for the images on the upper and lower rows, respectively. **B)** Whole-mount stained mammary glands shown in (A) are sectioned and imaged to better visualize the early stage tumors. Representative micrographs show that tumors grow as protrusions along primary ducts. Scale bars represent 1 mm for the left-most image and 0.1 mm for the high magnification images. **C)** Whole-mount staining results of the No:4 inguinal gland of an adult MMTVNIC; R26R-mT/mG mouse along with the micrographs for the endogenous expression of tdTomato and GFP demonstrate that the whole mammary ductal tree is formed by Neu^pos^ lineage cells despite the appearance of early-stage tumors only in few mammary ducts. Micrographs are representative of n=3 mice. Scale bars represent 1 cm and 1 mm for the images on the upper and lower rows, respectively.

The beads-on-string pattern of early-stage tumors being present in only a few mammary ducts in these glands may reflect the differences in the developmental history of each individual primary duct that diverges from each other via bifurcation during their pubertal development. Although the MMTV promoter provides a continuous overexpression of Neu oncogene throughout the whole mammary ductal tree during the entire postnatal life of these mice, the appearance of tumors only in some part of the mammary ductal tree with a relatively long latency of several months suggests that tumor initiation may require the accumulation of additional somatic mutations. Consequently, it is imperative that only those few mammary epithelial cells that have accumulated an adequate set of tumorigenic mutations—in addition to Neu overexpression—may generate tumors. Importantly, these cells would pass the acquired somatic mutations onto their progeny, which will also be located along the same primary/secondary ducts. This might provide distinct developmental histories of genomic mutations that hierarchically differ across the entire mammary ductal tree. Thus, we hypothesized that the beads-on-string pattern of tumorigenesis observed in MMTV-NIC glands might be the consequence of somatic mutagenesis occurring in a stochastic manner during pubertal stage of mammary gland development. To test this hypothesis, we analyzed early-stage tumor formation in MMTV-rTTa;TetON-HER2 double-transgenic females with an onset of HER2 overexpression at 15 weeks old adult stage, which is long after the completion of ductal elongation. Whole-mount staining of their No:4 glands demonstrated that, instead of displaying a beads-on-string pattern, early-stage multi-nodal tumors were randomly distributed within the mammary ductal tree (Supplementary Figure 2). However, it is also important to note that, although HER2 was overexpressed throughout the whole mammary ductal tree in these mice, tumors developed in only a few locations within the ductal tree, suggesting that HER2 overexpression alone is not sufficient for tumor initiation.

Taken together, our observations of early-stage tumorigenesis in these mouse models indicate that HER2/Neu-induced tumorigenesis is likely to require the accumulation of additional somatic mutations. Therefore, only those few mammary epithelial cells that have accumulated the adequate tumor-inducing set of mutations may give rise to mammary tumors. This proposed model could explain the observed beads-on-string pattern of tumorigenesis in MMTV-NIC glands and the relatively long latency of palpable tumor formation in both mouse models. Furthermore, as BCSCs arise from a Neu^neg^ cell-of-origin in MMTV-NIC tumors, these results may imply that tumor evolution could require the accumulation of somatic mutations by the cell-of-origin of BCSCs as a result of HER2/Neu overexpression in mammary epithelia.

### Neu overexpression results in ductal ectasia in mammary glands that contain poorly differentiated mammary epithelial cells

Most of previous studies using mouse models of HER2+ breast cancer focused on addressing the processes that took place in tumors rather than in mammary ducts prior to tumor formation^19, 20^. Thus, we sought to characterize the Neu overexpression-induced cellular and molecular changes in pre-cancerous mammary ducts. First, we performed whole-mount staining of mammary glands from MMTV-NIC mice at different ages to determine the oldest age, at which the No:4 glands could still be considered tumor-free (Supplementary Figure 3A). Inguinal glands obtained from some of the 10- and 15-week-old MMTV-NIC mice had early-stage tumors. In contrast, none of the No:4 glands from 6- or 8-week-old mice had macroscopically apparent tumors, even though they had ectatic duct structures on their primary ducts. These ductal ectasias appeared as tiny dense spots on primary ducts in whole-mount images (Figure 4A). Hematoxylin/Eosin-staining analysis revealed that these ectatic ducts are composed of multiple small acini-like protrusions along primary ducts (Figure 4B). A similar phenotype of ductal ectasia was previously reported in other mouse models of HER2+ breast cancer^15, 21^, even though a detailed cellular characterization was not documented.

**Figure 4.**
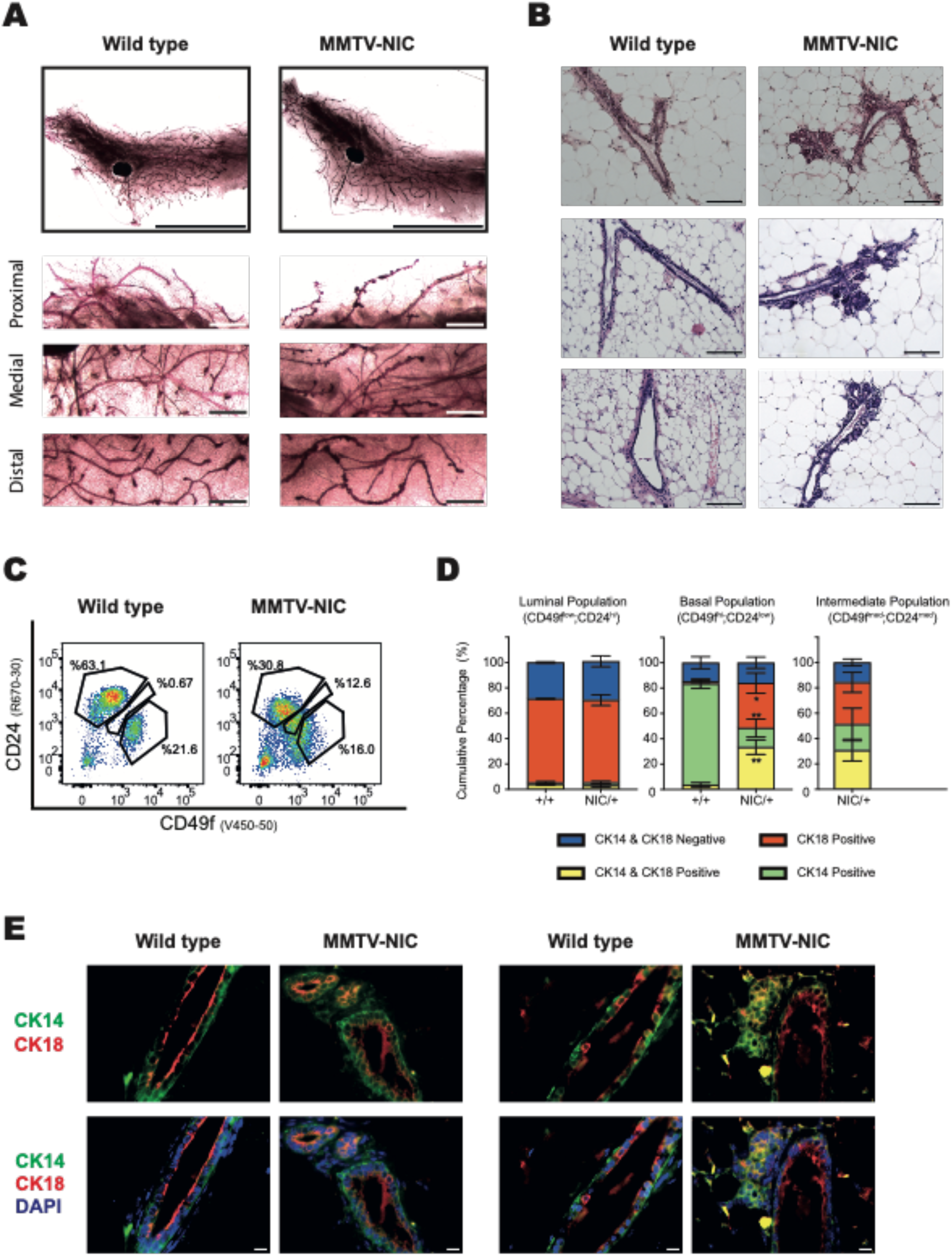
Neu overexpression results in ductal ectasia in mammary glands that contain poorly differentiated mammary epithelial cells. **A)** Whole-mount staining of the No:4 glands of 8-week-old MMTV-NIC and wildtype littermate mice shows the ectatic ducts as tiny dark spots along the primary ducts in Neu-overexpressing mammary glands. Higher magnification images from different parts of the glands are also shown. These images are representative of n=7 mice. Scale bars represent 1 cm and 1 mm for the low-magnification and high-magnification images, respectively. **B)** Hematoxylin/eosin staining of the microtome sections of No:4 glands of 8-week-old MMTV-NIC and wildtype littermate mice demonstrates their histological appearances. Micrographs shown are images from the glands of 2 different pairs of mice and are representative of n=10 mice. Scale bars represent 100 um. **C)** Flow cytometry analysis of No:4 mammary glands of 8-week-old MMTV-NIC and wildtype littermate mice shows an altered distribution of Lin^neg^ cells for their surface antigen profile of CD24 versus CD49f expression. Gates were set based on the populations present in the wildtype littermates. Dot plots shown are representative of n=4 pairs of mice. **D)** Co-immunofluorescence staining of primary mammary epithelial cells, which were sorted by using the gates shown in (**C**), for CK18 versus CK14 expression reveals an altered distribution of cell types within the basal and intermediate subpopulations. Values in the graph represent mean ± SEM of 3 pairs of mice and a total of 200 to 300 cells were counted for quantification purposes. *:p<0.05; **:p<0.01; by multiple unpaired t-test comparisons. **E)** Co-immunofluorescence staining of microtome sections of No:4 glands from 8-week-old MMTV-NIC and wildtype littermate mice for the expression of CK18 versus CK14 reveals that CK18^+^CK14^+^ cells are predominantly located in the ectatic ducts within MMTV-NIC glands. Shown are images from two different pairs of mice and are representative of n=3 pairs of mice. Scale bars represent 10 um.

Next, we performed flow cytometry analysis of primary cells obtained from No:4 glands of 8-week-old MMTV-NIC mice to determine whether Neu-overexpression may alter mammary epithelial lineages (Figure 4C). As expected, the surface antigen profiles for CD24 versus CD49f expression identified the luminal (CD24^hi^CD49f^lo^) and basal (CD24^lo^CD49f^hi^) epithelial lineages as distinct cell subpopulations in wildtype glands. In contrast, in MMTV-NIC glands the luminal and basal populations merged with the appearance of an intermediate population that could be described as (CD24^med^CD49f^med^). We then sorted these cells, cytospun on glass slides and stained with antibodies against the luminal cell marker CK18 and basal cell marker CK14 to determine the exact nature of their lineages (Figure 4C,D and Supplementary Figure 3B). In agreement with earlier studies, the CD24^hi^CD49f^lo^ and CD24^lo^CD49f^hi^ subpopulations from wildtype glands contained predominantly CK18^+^ luminal or CK14^+^ basal cells, respectively. The CD24^hi^CD49f^lo^ cell subpopulation from MMTV-NIC glands were also composed of predominantly CK18^+^ luminal cells similar to their wildtype littermates. However, both CD24^lo^CD49f^hi^ and CD24^med^CD49f^med^ subpopulations of MMTV-NIC epithelia displayed an abnormal composition of cells containing approximately 20% CK14^+^ basal, 30% CK18^+^ luminal, and 30% CK14^+^CK18^+^ cells. The mammary epithelial cells that express both luminal- and basal-specific cytokeratins are considered as the poorly differentiated epithelial cells and/or the bilineage progenitors^22–24^. To determine the location of these CK14^+^CK18^+^ cells in MMTV-NIC glands, we then performed immunostaining of mammary gland sections (Figure 4E). Although an inner layer of CK18+ luminal and an outer layer of CK14+ basal cells were observed in bilayered ductal structures of both wildtype and MMTV-NIC glands, the CK14^+^CK18^+^ epithelial cells were mostly confined to the ectatic ducts within MMTV-NIC glands. Importantly, these ectatic ducts were not composed of Ki67+ proliferative cells as is the case in the MMTV-NIC tumors (Supplementary Figure 3C,D), suggesting that these structures have not yet developed into a proliferative cancerous lesion. Furthermore, ectatic ducts did not contain an unusual number of apoptotic cells, marked with cleaved Caspase-3 staining (Supplementary Figure 3E), suggesting that they are not transient structures that undergo rapid involution.

Taken together, our results demonstrate that inguinal mammary glands of 8 weeks old MMTV-NIC female mice are free of tumors, despite containing ectatic duct structures, which are predominantly composed of poorly differentiated mammary epithelial cells.

### Neu overexpression results in altered metabolic activities, ROS accumulation and upregulation of PGP in mammary ducts

HER2 is a member of the EGFR family of receptor tyrosine kinases and has crucial functions in regulating tumor cell survival, proliferation, metabolism and invasiveness via modulating MAPK, PI3K, PKC, and STAT signaling^25–28^. To determine the molecular pathways altered in MMTV-Neu mammary ducts prior to tumor formation, we performed a proteomics analysis of intact mammary ducts isolated from 8-week-old MMTV-NIC mice and their wildtype littermates. Interestingly, there was only a single protein, the phosphoglycolate phosphotase (PGP), that showed a significant upregulation in MMTV-NIC ducts at the individual protein level (Figure 5A). To confirm this finding, we performed immunostaining for PGP on sections of MMTV-NIC and wildtype glands (Figure 5B). Our results demonstrated that ectatic duct structures within MMTV-NIC glands express an elevated level of PGP, while the bilayered mammary ducts in both MMTV-NIC and wildtype glands have lower levels of PGP expression. PGP is an evolutionary conserved metabolite repair enzyme that eliminates toxic by-products of glycolytic pathways, such as 2-phospho-L-Lactate and 4-Phosphoerythronate^29–31^. Furthermore, PGP can also dephosphorylate 3-Phosphoglycolate, a toxic byproduct of oxidative DNA damage repair enzymes such as tyrosyl DNA phosphodiesterase-1 (TDP1)^32–34^. Thus, the upregulation of PGP in the ectatic ducts within MMTV-NIC glands may suggest an alteration of metabolic activities and/or oxidative stress response.

**Figure 5.**
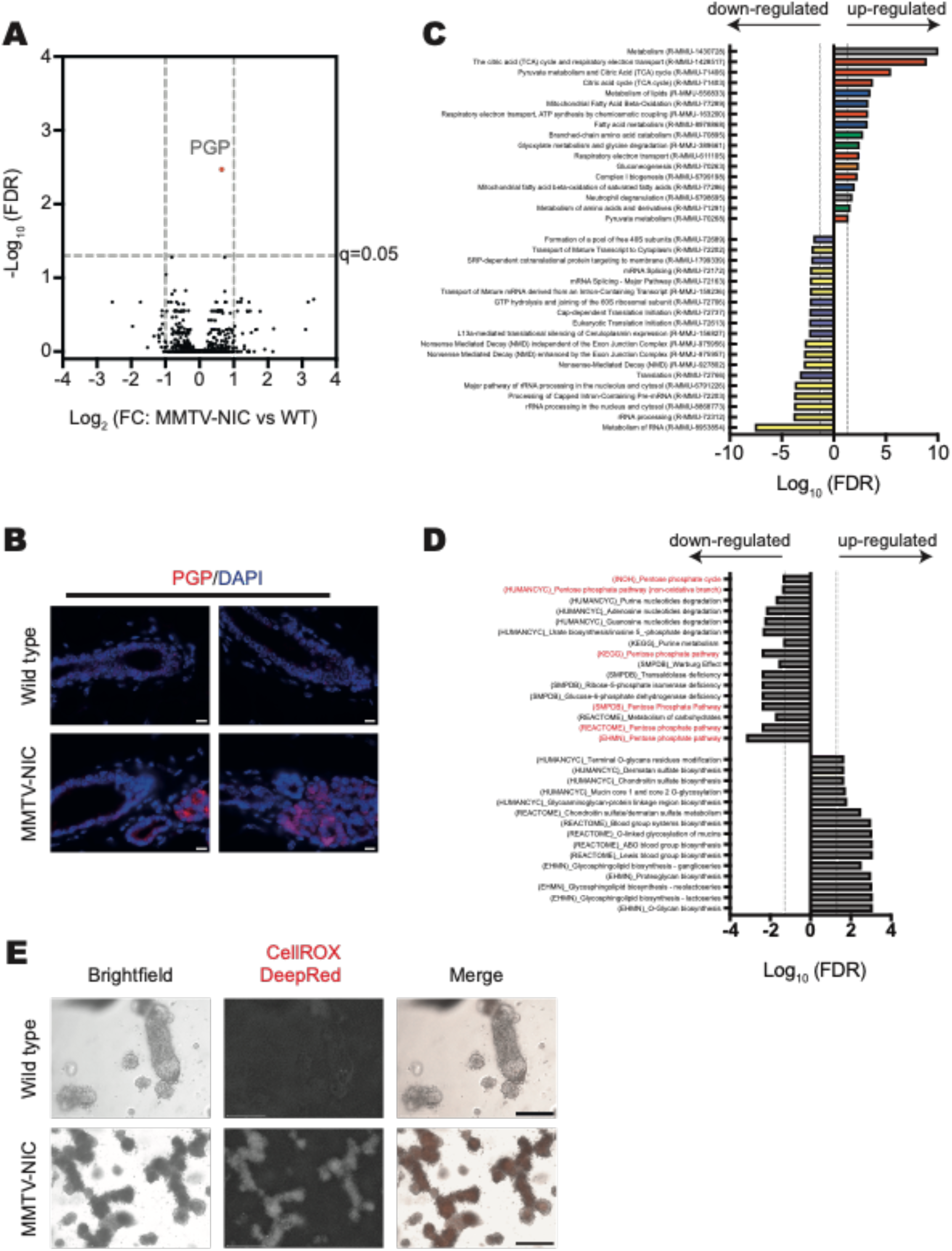
Neu overexpression results in altered metabolic activities, ROS accumulation and upregulation of PGP in pre-cancerous mammary ducts. **A)** Volcano plot analysis of differentially expressed proteins in intact mammary ducts of 8-week-old MMTV-NIC mice compared with their wildtype littermates. The only significant (q<0.05) alteration observed is PGP upregulation. (n=3 per genotype) **B)** Immunofluorescence staining of microtome sections of No:4 glands from 8-week-old MMTV-NIC and wildtype littermate mice reveals an upregulation of PGP levels in ectatic ducts within MMTV-NIC glands. Micrographs are representative of 3 pairs of mice analyzed. Scale bars represent 10 um. **C)** Reactome pathway enrichment analysis of the proteomics dataset for intact mammary ducts of 8-week-old MMTV-NIC mice compared with their wildtype littermates identifis significant upregulation of pathways related to glucose (in red) and lipid catabolism (in blue) along with significant downregulation of pathways related to RNA (in yellow) and protein anabolism (in purple). Vertical dashed lines represent the border of q=0.05. **D)** Pathway enrichment analysis of the metabolomics dataset for intact mammary ducts of 8-week-old MMTV-NIC mice compared with their wildtype littermates reveals the downregulation of the pentose phosphate pathway (in red) in MMTV-NIC ducts as a consistent finding from all 6 databases used. Th graph shows top 5 pathways identified in each database. The entire pathway enrichment dataset is shown in Supplementary Figure 4. Vertical dashed lines represent the border of q=0.05. **E)** CellROX Deep-Red staining of intact mammary ducts cultured in 3D Matrigel environment for 2 days demonstrates an increased accumulation of ROS in MMTV-NIC ducts compared to their wildtype controls. Micrographs are representative of n=3 mice per genotype. Scale bars represent 1 mm.

Next, we analyzed our proteomics dataset using Reactome pathway database to elucidate the molecular pathways altered in MMTV-NIC mammary ducts (Figure 5C). Interestingly, almost all significantly altered pathways were related to metabolism. Compared with wildtype controls, MMTV-NIC ducts have an upregulated proteome related to glucose and lipid catabolism and downregulated proteome of RNA and protein anabolism pathways. To further investigate how these changes may result in the alteration of metabolic activities in MMTV-NIC ducts, we performed an untargeted metabolomics analysis using mammary ducts isolated from 8-week-old mice and analyzed the annotated metabolites against 6 different pathway databases (Figure 5D and Supplementary Figure 4). All 6 database searches revealed a significant downregulation of the pentose-phosphate pathway activity in MMTV-NIC ducts compared with their wildtype controls.

The pentose-phosphate pathway is an anabolic pathway that runs parallel to glycolysis and is responsible for the production of (i) ribose-5-phosphate, an essential precursor for nucleotide synthesis; (ii) erythrose 4-phosphate, used for aromatic amino acid synthesis; and (iii) NADPH, which is used in reductive biosynthesis reactions, such as fatty acid synthesis, and acts as an anti-oxidant defense mechanism in cells by scavenging reactive oxygen species (ROS)^35, 36^. Thus, the downregulation of the pentose-phosphate pathway activity might contribute to the observed downregulation of RNA and protein anabolisms in MMTV-NIC ducts. Furthermore, since glycolysis and pentose-phosphate pathways compete for the use of glucose-6-phosphate, the downregulation of pentose-phosphate pathway activity may allow increased glycolysis and subsequently the citric acid cycle, as observed in our proteomics dataset for MMTV-NIC ducts. Given that the cellular energy production involving both glucose and lipid catabolism are the main sources of ROS production, while the main contributor of the ROS-scavenging NADPH is the pentose-phosphate pathway in mammalian cells, we hypothesized that the observed alterations in the metabolic pathways within MMTV-NIC ducts may result in ROS accumulation. To test this hypothesis, we employed a 3D *ex vivo* organ culture model to detect ROS accumulation, as it is not plausible to visualize ROS in fixed tissue samples. Intact mammary ducts isolated from inguinal glands of MMTV-NIC mice and their wildtype littermates were plated in 3D-Matrigel culture and stained with CellROX Deep Red reagent two days after plating. Our results demonstrated a visually obvious increase in CellROX staining in MMTV-NIC ducts compared to wildtype ducts (Figure 5E), suggesting that Neu overexpression in mammary epithelia results in ROS accumulation. To address whether the ROS accumulation is a direct effect of HER2/Neu overexpression rather than the consequence of the architectural changes within MMTV-NIC glands, we generated a transgenic murine mammary epithelial cell line, EPH4, with doxycycline-inducible overexpression of *ERBB2/HER2* (Supplementary Figure 5). Induction of HER2 overexpression in these cells led to ROS accumulation (Supplementary Figure 5C), indicating that this phenotype is a direct consequence of HER2/Neu overexpression.

Collectively, our results demonstrate that Neu overexpression in the mammary gland results in significant alterations of metabolic pathways including glucose, lipid, RNA, and protein metabolisms. As the activities of these metabolic pathways are interconnected, the integrated network analysis for our proteomics and metabolomics results suggests that the downregulation of pentose-phosphate pathway activity in MMTV-NIC ducts might be central to most of these metabolic changes (Supplementary Figure 6). Furthermore, we observed increased ROS accumulation in HER2/Neu-overexpressing mammary epithelia and upregulation of PGP levels within the ectatic ducts of MMTV-NIC glands. This may suggest that altered energy metabolism in MMTV-NIC ducts may result in oxidative stress that can lead to oxidative DNA damage.

### Neu overexpression in mammary glands results in the upregulation of oxidative stress and DNA damage markers leading to the accumulation of somatic mutations in both Neu^pos^ and Neu^neg^ lineages

Low levels of ROS regulate cellular and extracellular signaling pathways and thus have an impact on cell metabolism, proliferation, and differentiation^37^. Most types of ROS can diffuse through cell membrane, modify extracellular matrix proteins and act as a signaling molecule on the neighboring cells^38^. However, at high levels, ROS can lead to oxidative stress in the tissue microenvironment with genotoxic effects by inducing DNA damage and other deleterious effects by damaging lipids and proteins^39–41^.

To determine whether increased ROS levels in MMTV-NIC mammary ducts lead to the upregulation of oxidative stress markers in the tissue microenvironment, we analyzed the glands from 8 weeks-old MMTV-NIC/+;R26R-mT/mG mice and their wildtype littermates via immunostaining for oxidative stress markers 4-HNE and 8-oxoGuanine along with the anti-GFP antibody to distinguish the Neu^pos^ and Neu^neg^ lineages. The lipid peroxidation marker 4-HNE was upregulated in MMTV-NIC/+;R26R-mT/mG glands and observed to be present in both Neu^neg^ and Neu^pos^ lineage cells (Figure 6A), suggesting that increased ROS levels also affects those cells that do not express the Neu oncogene themselves. Similarly, the oxidative DNA damage marker 8-oxoGuanine was observed in the nuclei of both the Neu^neg^ and Neu^pos^ lineage cells within the ectatic ducts of these glands (Figure 6B), indicating the oxidative DNA damage being present also in the neighboring cells of Neu^pos^ lineage cells. Previous studies showed that high levels of oxidative stress may lead to double-strand DNA breaks^42^. Thus, we analyzed Gamma-H2Ax levels in MMTV-NIC/+;R26R-mT/mG glands and demonstrated the presence of Gamma-H2Ax staining in a few cells within the ectatic ducts, including some of the Neu^neg^ lineage cells (Figure 6C).

**Figure 6.**
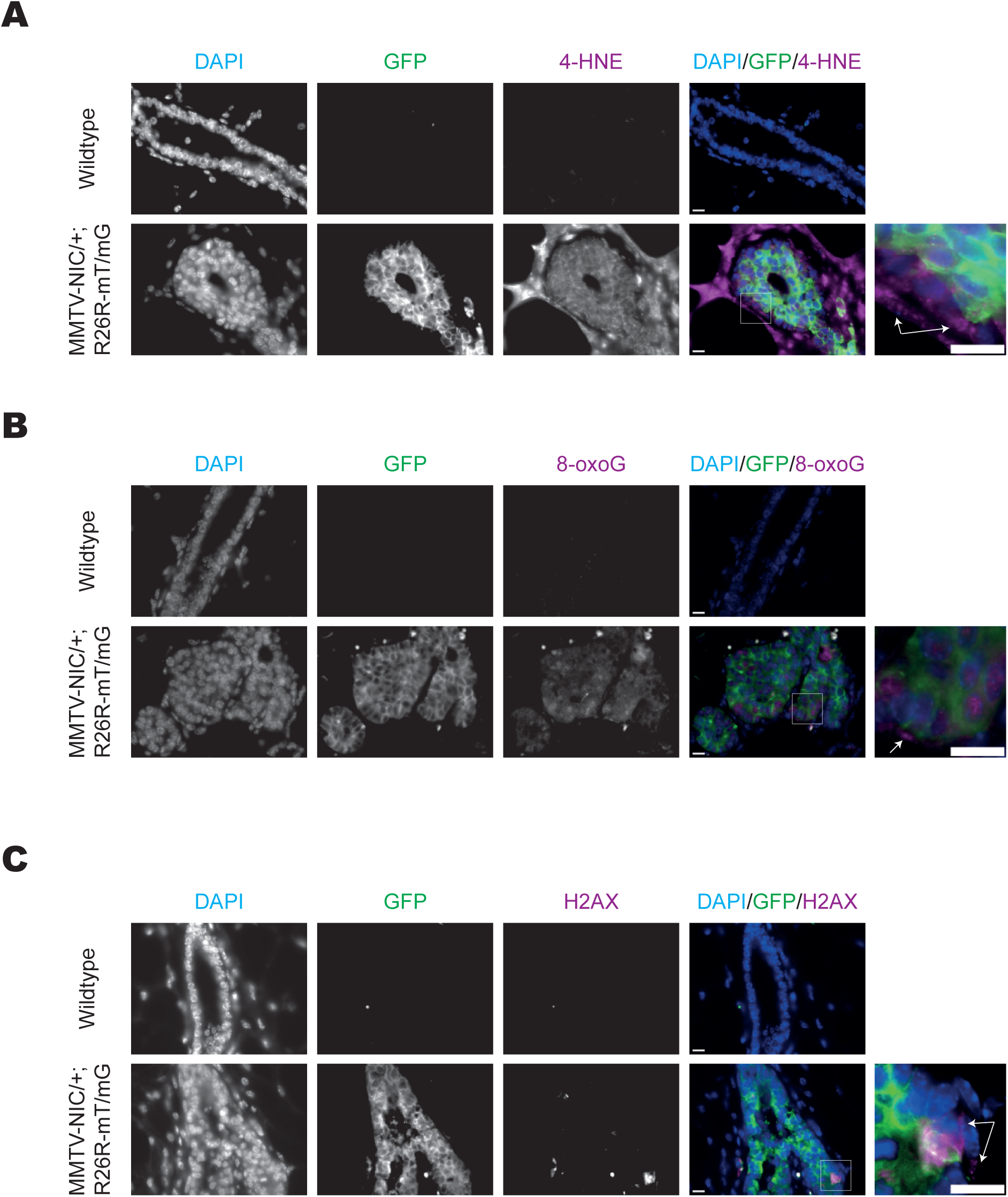
Neu overexpression leads to the upregulation of oxidative stress and DNA damage markers in both Neu^pos^ and Neu^neg^ lineages in pre-cancerous mammary glands. **A-C)** Co-immunofluorescence staining of microtome sections of No:4 glands from 8-week-old MMTV-NIC;R26R-mT/mG and wildtype littermate mice for the expression of GFP -marker of the Neu^pos^ lineage-versus 4-HNE (**A**), 8-oxoGuanine (**B**), and gamma H2AX (**C**) reveals an upregulation of all these markers in both Neu^pos^ and Neu^neg^ lineage cells. Arrows in the high-magnification images point to the Neu^neg^ lineage cells expressing the corresponding stress/damage markers. Micrographs shown are representative of 3 pairs of mice analyzed. Scale bars represent 10 um.

Next, we investigated whether the observed PGP upregulation in ectatic ducts could be a result of the oxidative stress. The immunostaining of MMTV-NIC/+;R26R-mT/mG mammary gland sections for PGP demonstrated slightly different patterns in their ectatic ducts versus bilayered ductal structures (Supplementary Figure 7). PGP expression was present only in the Neu^pos^ lineage cells within the bilayered ductal structures, whereas both Neu^pos^ and Neu^neg^ lineage cells expressed PGP in ectatic ducts. This may reflect two distinct functions of PGP, namely the regulation of glucose metabolism and the oxidative stress response, and therefore the higher levels of PGP in ectatic ducts could be due to the increased levels of oxidative DNA damage within those cells.

Given that DNA damage does not necessarily result in mutagenesis, we sought to determine whether Neu^neg^ lineage cells within MMTV-NIC/+;R26R-mT/mG tumors indeed acquire any somatic mutations during tumor evolution. To this end, we performed whole genome sequencing analysis of tdTOM^+^ and GFP^+^ tumor cells and analyzed their somatic mutation signatures. The overall frequency of single nucleotide variants (SNVs) and InDels was similar between tdTOM^+^ and GFP^+^ tumor cells (Supplementary Figure 8), indicating that the mutation burden in Neu^neg^ lineage cells is similar as in Neu^pos^ lineage cells. Importantly, the observed mutation signatures were not only similar between the tdTOM^+^ and GFP^+^ tumor cells within each tumor, but also between different tumors (Figure 7A,B). This may suggest that tumor evolution in different tumors may depend on similar etiologies. Thus, we evaluated these mutation signatures by analyzing the cosine similarity to existing mutational signatures observed in a large set of human cancers^43^ to determine whether there may be a common etiology involved in tumor initiation and progression. For the SNVs, the most common profile observed was SBS-5 (Figure7C), a “clock-like’ profile indicating that the number of mutations correlate with age. Interestingly, SBS-5 is also one of the six most common profiles found in breast tumors in women^44^. For the InDels, the ID1 and ID2 profiles that are potentially linked to mismatch repair deficiencies, and ID12 profile with unknown etiology were detected (Figure 7D). Although ROS-induced DNA damage is mainly repaired by the base-excision and nucleotide-excision repair pathways, the mismatch repair pathway has also been implicated in oxidative DNA damage response, particularly for the 8-oxoG residues^45^. Therefore, our results indicate that somatic mutagenesis occurring during tumor evolution can also be observed in Neu^neg^ lineage cells through the same etiologies as the Neu^pos^ lineage cells and are likely caused by oxidative stress and developmental clock involved in tumorigenesis.

**Figure 7.**
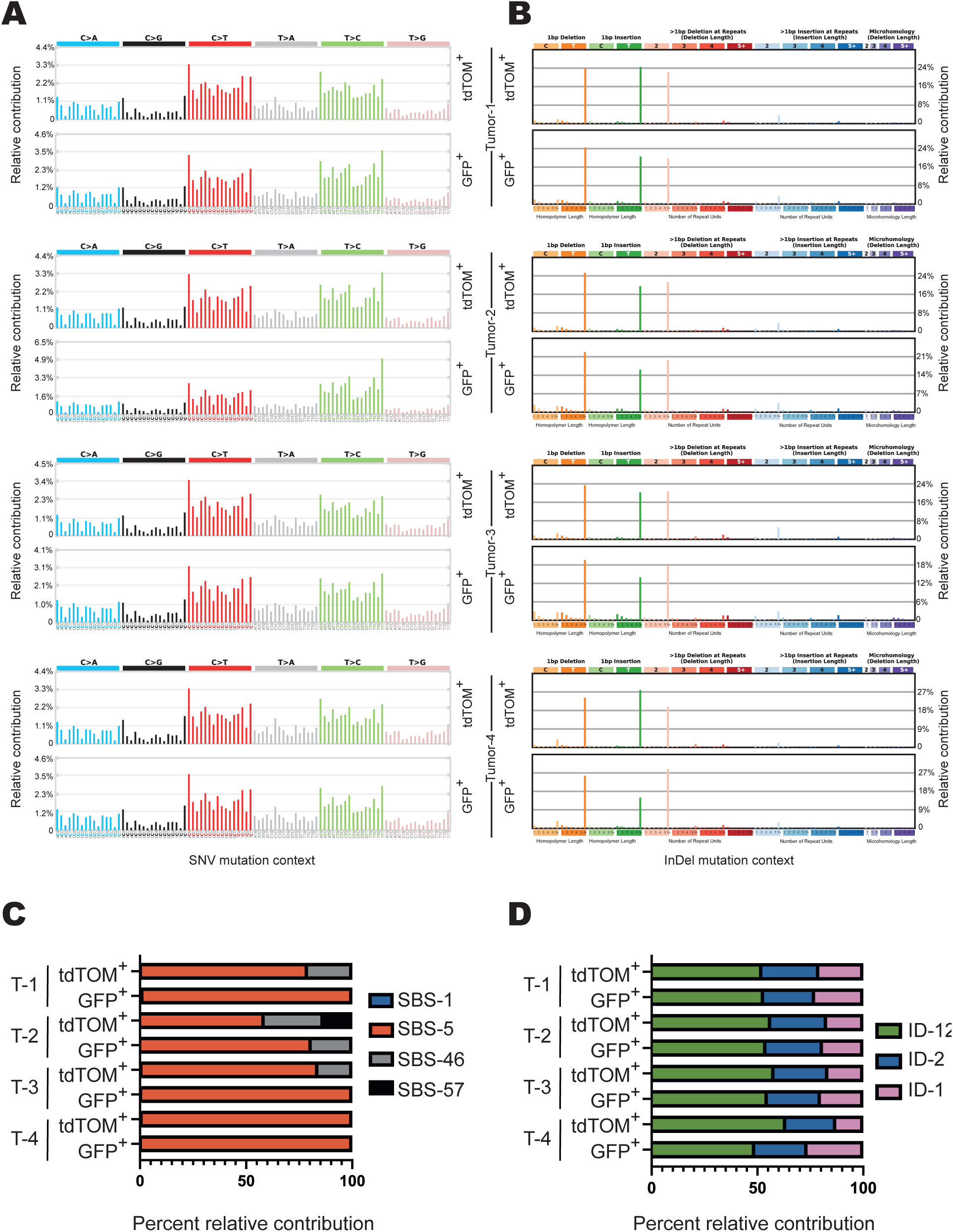
Neu overexpression driven mammary tumorigenesis results in the accumulation of somatic mutations in both Neu^pos^ and Neu^neg^ lineage tumor cells. **A,B)** Somatic SNV (**A**) and InDel (**B**) mutation signatures of tdTOM^+^ Neu^pos^ and GFP^+^ Neu^neg^ lineage cells for 4 independent tumors showed highly similar profiles. Note that the potential germline mutations has been subtracted from the dataset during analyses using the whole genome sequencing data obtained for tail biopsy samples of each corresponding tumor-bearing MMTV-NIC;R26R-mT/mG mice. **C,D)** Deconvolution of the SNV **(C)** and InDel **(D)** mutation signatures of tdTOM^+^ Neu^pos^ and GFP^+^ Neu^neg^ lineage cells for 4 independent tumors into known COSMIC signatures found in human tumors identified represented profiles as SBS-1, SBS-5, SBS-46, SBS57, ID-12, ID-2, and ID-1. T: Tumor.

Taken altogether, our results demonstrate that Neu overexpression in mammary ducts alters the metabolism, which leads to increased levels of genotoxic oxidative stress in their tissue microenvironment. Consequently, this leads to DNA damage and accumulation of somatic mutations in both Neu^pos^ and Neu^neg^ lineage cells, the latter of which contains the cells-of-origin of BCSCs. However, as these somatic mutations are random in their nature, only potential cells-of-origin of BCSCs that have acquired an adequate tumor-inducing set of somatic mutations will give rise to tumors. This proposed model may explain the observed ductal lineage dependency of tumorigenesis in MMTV-NIC mammary glands and illustrate a previously unrecognized non-cell-autonomous mechanism of HER2/Neu-induced breast tumorigenesis during the earliest tumor initiation stages.

## DISCUSSION

Previous studies revealed the crucial role of HER2 in regulating the mitogenic signaling pathways important for tumor progression in HER2+ breast cancer, leading to the advent of successful HER2-targeting treatments^25–28^. Furthermore, various mouse models of HER2+ breast cancer demonstrated that HER2 is a driver oncogene for breast cancer, as female mice with transgenic HER2/Neu overexpression in mammary glands develop tumors with 100% prevalence^19, 20^. However, to date it is still not well understood at a cellular level how HER2/Neu overexpression leads to tumor initiation *in vivo*. Here, we have shown that Neu overexpression in mammary epithelia alters their metabolism, leading to ROS accumulation in the tissue microenvironment due to the upregulation of glucose and lipid catabolism together with the downregulation of pentose-phosphate pathway. Increased ROS causes upregulation of oxidative stress and DNA damage markers, and eventually leads to the accumulation of somatic mutations in both Neu^pos^ and Neu^neg^ lineage cells. Using clinical samples of DCIS and two distinct mouse models, we have shown that BCSCs and their cells-of-origin do not express HER2/Neu oncogene. Taken together, our results illustrate that HER2/Neu overexpression non-cell-autonomously induces mutagenesis in the surrounding HER2/Neu-negative cells that contain the cells-of-origin of BCSCs. As a result of the stochastic nature of the somatic mutagenesis, tumor initiation may require the accumulation of an adequate tumor-inducing set of mutations that can be acquired by only a few cells. Consequently, this may explain why, although HER2/Neu is overexpressed throughout the ductal tree in our mouse models, early-stage tumors form with a relatively long latency and in only a few mammary ducts.

The cell type for the cell-of-origin of BCSCs was addressed by several studies, though a consensus has not yet been established^46^. Candidates include the multi-potent mammary epithelial stem cells (MaSCs), unipotent mammary stem cells, luminal progenitors, or parity-induced mammary epithelial cells (PI-MECs). In the MMTV-Neu mouse model, our results demonstrate that the cell-of-origin of BCSCs reside within the Neu^neg^ lineage, even though we cannot propose a specific cell type as the likely cell-of-origin. However, given that tumor initiation requires time to acquire an adequate set of tumor-inducing somatic mutations, and the early-stage mammary tumors display a ductal lineage dependency, it is imperative that the cell-of-origin of BCSCs must be a long-living cell with self-renewal capacity that is functionally involved in ductal elongation processes during mammary gland development. Furthermore, BCSCs are a heterogenous population of cells known for their plasticity, as non-stem tumor cells may also de-differentiate into BCSCs under certain circumstances, such as cytoablative treatments^6^. Our lineage-tracing experiments showed that tdTOM^+^ Neu^neg^ lineage cells formed mammospheres *ex vivo* and initiated tumors upon transplantation *in vivo*, as expected for a population that contains BCSCs. In contrast GFP^+^ Neu^pos^ lineage cells did not contain mammosphere-forming BCSCs, although they were still able to initiate tumors when transplanted into mammary fat pads. Interestingly, mammosphere-forming BCSCs could be recovered from tumors that were formed by the GFP^+^ Neu^pos^ lineage cells upon transplantation into the fat pad (data not shown). This observation suggests that some of the non-stem cells within Neu^pos^ lineage tumor cells might have de-differentiated into BCSCs either due to the dissociation into single cells for cell sorting or due to the sudden microenvironmental change of being injected into the fat pad, a situation that resembles to the transition from DCIS into invasive breast cancer when early-stage tumor cells break through the ductal basal lamina and invade the mammary stroma. Future studies will be needed to address this type of BCSC plasticity that may occur during the malignant transformation of breast tumors.

DCIS is usually asymptomatic and most commonly found through breast screening as microcalcifications on mammograms. Interestingly, a beads-on-string pattern of microcalcifications were observed in the mammograms of some patients diagnosed with DCIS^47^. Thus, our findings in the MMTV-NIC mouse line for the ductal lineage-dependency of early-stage tumors may also have clinical relevance to explain the tumor evolution of premalignant stage breast cancer in women. Given that both ectatic ducts and early-stage tumors form as protrusions from primary ducts in MMTV-NIC glands, it is likely that these ectatic ducts could serve as early-stage tumor precursors in this mouse model. Intriguingly, these ectatic ducts contain a considerable amount of poorly differentiated CK14^+^CK18^+^ epithelial cells, resembling the bipotent progenitors or MaSCs of the embryonic mammary glands^23, 24^. Hence, it would be necessary to determine in future studies whether these poorly differentiated cells within ectatic ducts could be the cells-of-origin of BCSCs and whether they also exist in atypical ductal hyperplasia lesions in clinical samples.

Current clinical practice does not require a pathological assessment of HER2 status in DCIS, as its status does not influence the choice of treatment. Nevertheless, earlier studies showed that up to 70 % of high-grade DCIS are HER2+^3^. We also observed that 10 out of 15 DCIS samples we analyzed had more than 10% of DCIS cells expressing HER2 (Supplementary Figure 1). According to the mechanism we propose here, HER2 overexpression plays a non-cell-autonomous role in pre-malignancy; therefore, it is plausible that, after initial mutagenesis, some of the HER2+ DCIS transform into invasive tumors of other molecular subtypes depending on which specific mutations were acquired by BCSCs during the DCIS stage. This may provide an intriguing explanation for the observed discrepancy between the incidence of HER2+ DCIS and the much lower incidence of HER2+ invasive breast cancer. Finally, this study may prompt investigations into whether HER2-targeting agents with known bystander effects^48^ could be used to target the evolving BCSCs and their cells-of-origin as a preventive treatment in patients with DCIS.

## ACKNOWLEDGEMENTS

This work was supported by scientific fellowship grant (#2016MaySF722) to A.U from Breast Cancer Now (BCN). S.B.G was supported by Study Abroad Program (MEBYLSY2018) of the Ministry of National Education of the Republic of Türkiye. We wish to thank the Bioimaging, Histology, Bio-MS, Genomic Technologies, and Flow Cytometry facilities of Manchester University for core research facility support; staff at the Biological Safety Unit for looking after the mice; Olga Ucar for proofreading of the manuscript; and Dr Jean Zhao for kindly providing us with the MMTV-rTTa;TetON-HER2 mouse line.

The authors wish to acknowledge the roles of the BCN Tissue Bank in collecting and making available the samples and data, and the patients who have generously donated their tissues and shared their data to be used in the generation of this publication. We particularly thank Iain Goulding and Jenny Gomm for the preparation of ductal organoid samples and making them available for our study.

## AUTHOR CONTRIBUTIONS

S.B.G and A.U developed the concept of this study, analyzed data, and wrote the manuscript. K.J.W, A.G., and K.B. contributed to study design and data interpretation. O.W., F.C., S.B.G., and K.P. performed cytospin analyses; S.B.G., O.W., L.D., and A.U. performed histological analyses; S.B.G., F.C., and A.U. performed flow cytometry analyses; S.B.G., F.C., D.N., and A.U. performed mammosphere assays; S.B.G. and B.A.T. performed transplantation experiments; S.B.G., W.W., R.P., G.T., Y.O., and I.J.D. performed omics and WGS studies and bioinformatical analyses. All authors critically revised the manuscript and approved the final version.

## MATERIALS and METHODS

### Mouse experiments

All mouse experiments were conducted under project license in accordance with the UK Home Office Animals (Scientific Procedures) Act (1986) regulations and with the approval of study protocols by the Animal Welfare and Ethical Review Body (AWERB) of the University of Manchester. Mice were maintained in a pathogen-free facility and kept in 12-hours light-dark cycles in temperature- and humidity-controlled environment at the Biological Safety Unit of the University of Manchester and were provided with food and water ad libitum.

The mouse lines MMTV-NIC, R26R-mT/mG, and MMTV-rTTa;TetON-HER2 were previously described^15, 16^. All mouse lines were maintained in pure FVB/J background through continuous backcrossing with wildtype FVB/J mice. For the induction of HER2 overexpression in MMTV-rTTa;TetON-HER2 mice, female mice received food pellets containing doxycycline (625 mg/kg diet-Envigo #TD.120769).

For orthotopic transplantation experiments, adult (age range: 14-17 weeks; weight range: 22-26.5 gr) female nude mice in a CBA background were used as hosts. Cells sorted by FACS were resuspended in serum-free medium and mixed with Matrigel in a 1:1 ratio to achieve a cell concentration of 1X10^6^ cells per ml. Fifty microliter of these cell suspensions (50,000 cells) were injected into the mammary fat pad near the nipple area of either right or left No:4 mammary glands of mice under anesthesia. Transplanted mice were routinely monitored for tumor formation after the operation and culled when tumors reach 1 cm^3^ size.

### Primary cell isolation and mammosphere culture

Clinical DCIS samples were obtained from the Breast Cancer Now Tissue Bank as cryopreserved ductal organoids. After thawing, ductal organoids were resuspended and kept in mammosphere culture medium at 37 °C with 5% CO2 for 2 hours for acclimatization to improve cell survival. Next, they were digested using Trypsin-EDTA (Sigma) followed by digestion in 1 mg/ml Dispase II (#4942078001, Roche) and 0.4 mg/ml DNase I (#11284932001, Roche) to obtain single cell suspensions.

Primary cells from murine mammary tumors or murine mammary glands were isolated using the protocol described previously^18^. Mammosphere formation assay was performed as previously described in detail elsewhere^18, 49^.

### Flow cytometry and FACS sorting

Single cell suspensions were blocked with CD16/CD32 (#553141, Becton Dickinson (BD)) in FACS Buffer (2% FBS in PBS) and stained with primary antibodies, followed by staining with secondary antibodies. The dead/live cell markers used were either DRAQ7 (Biostatus), SytoxGreen (Invitrogen), DAPI (Sigma) or PI (Sisgma). Antibodies used for murine samples were anti-mouse CD45-Biotin (#553077, BD), anti-mouse CD31-Biotin (#558737, BD), anti-mouse Ter119-Biotin (#553672, BD), anti-mouse CD24-APC (#562349, BD), anti-mouse CD49f-eFluor450 (#48-0495-92, Invitrogen), anti-human CD340 (HER2) (#562349, BD), Streptavidin-PE-Cy7 (#557598, BD), or Streptavidin-FITC (#554060, BD) antibodies. Antibodies used for human samples were anti-human CD31 (#17031942, Invitrogen), anti-human CD45 (#17945942, Invitrogen), anti-human CD235a (#17998742, Invitrogen), anti-human CD326 (EpCAM) (#12932642, Invitrogen), and anti-human CD340 (HER2) (#562349, BD). Antibodies used for cell lines were anti-human CD340 (HER2) (#562349, BD). All flow cytometry experiments were performed using either LSR Fortessa or ARIA Fusion (BD), and data analyses were performed using FlowJo.

### Whole mount staining and imaging of mammary glands

Whole mount staining of mammary glands with Carmine-Alum was performed as previously described^50^. For the visualization of the endogenous tdTomato and GFP fluorescence as well as the brightfield imaging of whole Carmine-Alum-stained glands, a Leica M205 FA upright stereomicroscope was used.

### Histological analyses

Inguinal mammary glands dissected form mice were fixed in 4% PFA overnight, dehydrated in increasing concentrations of ethanol and embedded in paraffin wax using standard protocols. Five um thick sections were prepared using the Leica RM2255 microtome and collected on microscope slides. Sections were heated at 60°C for 30 minutes, deparaffinised with xylene, and rehydrated with incubations in decreasing concentrations of ethanol.

For haematoxylin/eosin staining, rehydrated sections were stained with haematoxylin (Sigma, HHS16) for 8 minutes, washed in tap water, acidified within 0.1% HCl in 70% ethanol, and washed in tap and deionized water. Sections were then stained with Eosin Y solution (Sigma #318906) activated by glacial acetic acid and washed with deionized water. After dehydration, sections were mounted and imaged using EVOS cell imaging system (Invitrogen) and analyzed using *Fiji ImageJ*.

For immunostaining on sections, rehydrated sections were boiled in 0.01M Citrate buffer for antigen retrieval, incubated in 0.05M Tris buffer and blocked with 10% horse serum in PBS at room temperature for 1 hour. For staining with mouse antibodies, additional blocking using the mouse-on-mouse (M.O.M) kit was used according to the manufacturer’s protocols. After blocking, sections were incubated with primary antibodies in blocking buffer at 4°C overnight. Sections were then washed and incubated in secondary antibodies in blocking buffer in dark at room temperature for 1 hour. DAPI (Sigma) was included in secondary antibody solutions for counterstaining of nuclei. Sections were then washed, dried and mounted. Images were collected on a Axio Imager.D2 Upright or Axio Imager.M2 Upright Zeiss microscope using a *63x / 1.4* Plan Apochromat (Oil; DIC) objective and captured using a Coolsnap HQ2 camera (Photometrics) through Micromanager software v1.4.23. Images were then processed and analysed using *Fiji ImageJ.* Primary antibodies used were: anti-Phosphoglycolate phosphotase (PGP) (ab242104, Abcam), anti-4 Hydroxynonenal (4HNE) (ab46545, Abcam), anti-8-oxoguanine (MAB3560, Merck), anti-phospho-histone H2AX (Ser 139) (3748823, Merck), anti-GFP (46825, Rockland), anti-GFP (A11122, Invitrogen), anti-Cytokeratin-14 (PA5-16722, Invitrogen), anti-Cytokeratin-18 (61028, Progen), anti-cleaved caspase 3 (#9661, Cell Signalling Technology), and anti-Ki67 (MA5-14520, Invitrogen). Secondary antibodies used were: Donkey anti-rabbit AlexaFluor 488 (#A21206, Invitrogen), donkey anti-mouse AlexaFluor 594 (#A21203, Invitrogen), donkey anti-rabbit AlexaFluor 647 (#A31573, Invitrogen), donkey anti-goat AlexaFluor 488 (#A11055, Invitrogen), donkey anti-rabbit AlexaFluor 555 (#A32794, Invitrogen), donkey anti-rabbit AlexaFluor 594 (#A21207, Invitrogen), and donkey anti-mouse AlexaFluor 647 (#A31571, Invitrogen).

### Cytospin and immunofluorescence

Primary cells sorted by FACS were cytospun on microscope slides and stained with antibodies as previously described^18^. Primary antibodies used were Anti-Cytokeratin-14 (PA5-16722, Invitrogen) and Anti-Cytokeratin-18 (61028, Progen); with secondary antibodies Donkey Anti-Rabbit AlexaFluor 488 (#A21206, Invitrogen) and Donkey Anti-Mouse AlexaFluor 594 (#A21203, Invitrogen).

### Intact mammary duct isolation and visualization of ROS

No:4 mammary glands were dissected and spread on sterile microscope slides. Lymph nodes were removed, and the glands were cut into thin horizontal stripes before digesting them with Collagenase A (#11088793001, Roche) at 37°C for 3–4 hours. The ductal organoids were then separated from single cells using a 100-um mesh and collected into a tube by flushing them with medium. Isolated intact ducts were mixed with Matrigel (#354234, Corning) to plate them as 3D culture in 12-well plates and cultured with mammosphere medium. After two days of culture, the accumulation of ROS in ductal organoids were visualized by using the CellROX Deep Red staining kit (Invitrogen) and imaged with EVOS cell imaging system (Invitrogen)

### Proteomics analysis of intact mammary ducts

Intact mammary ducts were isolated from No:4 mammary glands of 8-week-old mice using the protocol described above. Ductal organoid samples were then resuspended in 1X S-Trap lysis buffer and homogenized by sonication using a LE220-Plus focused ultrasonicator (Covaris) at 300 peak power, 100 cycles per burst and a 40% duty factor. Protein concentration was quantified using a BCA assay or by Millipore Direct Detect. Fifty ug of protein was reduced and alkylated using final concentrations of 5mM of Dithiothreitol (DTT, BP172-5, Fisher) and 15mM iodoacetamide (IAM, I1149, Sigma Aldrich). Proteins were then fixed to S-Trap spin columns (Protifi) using S-Trap binding buffer, and then eluted with 50mM TEAB, 0.1% formic acid (FA), and 30% acetonitrile containing 0.1% FA. Peptides were desalted for LC-MS/MS using POROS Oligo R3 beads (Thermo Fisher) in Corning FiltrEX desalt filter plates. Samples were analyzed by LC-MS/MS using UltiMate 3000 Rapid Separation LC system (RSLC, Dionex Corporation), coupled to a Q Exative HFTM mass spectrometer (Thermo Fisher). Features were identified and quantified using MaxQuant (version 1.6.14.0) and were searched against the Uniprot *Mus musculus* reference proteome (UP000000589, accessed 5th Sept 2020), including Oxidation (M)(P), Acetyl (protein N-term) as variable modification and Carbamidomethyl (C) as a fixed modification. Up to two missed cleavages were allowed. MS/MS was set to 8 ppm, while peptide tolerance was 0.015 Da. Peptide data was then processed in MSqRob, normalizing intensity values between samples via centre median. Mouse genotype (i.e. WT or MMTV-Neu) was set as fixed effect, while sequence and biological replicate were defined as random effects. A statistical enrichment test was performed with PANTHER Enrichment Test (released 28/07/2020) using Reactome pathway annotations (v.65), and applying False Discovery Rate correction. Pathways with q<0.05 were deemed significantly enriched. Pathway network data was visualized by mapping PANTHER derived values onto a custom Reactome network facsimile in Cytoscape (Open source, v.3.8.0).

### Metabolomics analysis of intact mammary ducts

Intact mammary ducts were isolated from No:4 glands of 8-week-old mice as described above. Ductal organoids were centrifuged, tissue pellets were snap-frozen in liquid nitrogen and then resuspended in 50% methanol to incubate overnight at 4 °C. Next day, samples were centrifuged at 20,000 x g for 3 min and the top 50 ul supernatant was transferred to a glass autosampler vial with 300 µl insert and capped. Quality control samples were made by pooling 10 µl from each sample.

Liquid chromatography-mass spectrometry analysis was performed using a Thermo-Fisher Ultimate 3000 HPLC system consisting of an HPG-3400RS high pressure gradient pump, TCC 3000SD column compartment and WPS 3000 Autosampler, coupled to a SCIEX 6600 TripleTOF Q-TOF mass spectrometer with TurboV ion source. The system was controlled by SCIEX Analyst 1.7.1, DCMS Link and Chromeleon Xpress software.

A sample volume of 5 μL was injected by pulled loop onto a 5 μL sample loop with 150 μl post-injection needle wash with 9:1 acetonitrile and water. Injection cycle time was 1 min per sample. Separations were performed using an Agilent Poroshell 120 HILIC-Z column with dimensions of 150 mm length, 2.1 mm diameter and 2.7 μm particle size equipped with a guard column of the same phase. Mobile phase A was water with 10 mM ammonium acetate adjusted to pH 9 with ammonium hydroxide and 20 µM medronic acid, mobile phase B was 85:15 acetonitrile and water with 10 mM ammonium acetate adjusted to pH 9 with ammonium hydroxide and 20 µM medronic acid. Separation was performed by gradient chromatography at a flow rate of 0.25 ml/min, starting at 96 % B for 2 minutes, ramping to 65 % B over 20 min, hold at 65 % B for 2 min, then back to 96 % B. Re-equilibration time was 5 min. Total run time including 1 min injection cycle was 30 min. The mass spectrometer ran in negative mode under the following source conditions: curtain gas pressure, 50 psi; ionspray voltage, -4500 V; temperature, 400 °C; ESI nebulizer gas pressure, 50 psi; heater gas pressure, -70 psi; declustering potential, -80 V. Data was acquired in an information dependent manner across 10 high sensitivity product ion scans, each with an accumulation time of 100 ms and a TOF survey scan with accumulation time of 250 ms. Total cycle time was 1.3 s. Collision energy was determined using the formula CE (V) = 0.084 x m/z +12 up to a maximum of 55 V. Isotopes within 4 Da were excluded from the scan.

Acquired data were checked in PeakView 2.2 and imported into Progenesis Qi 2.4 for metabolomics, where they were aligned, peaks were picked, normalized to all compounds and deconvoluted according to standard Progenesis workflows. Annotations were made by searching the accurate mass, MS/MS spectrum and isotope distribution ratios of acquired data against the NIST MS/MS metabolite library. Metabolites were identified by searching retention times and accurate masses against an in-house chemical standard library. A validated identification is only given if identical hits are made against both the NIST MS/MS and in-house chemical standard libraries. Samples were grouped according to the mouse genotypes (i.e WT or MMTV-Neu) and were filtered based on the following statistical criteria – a maximum fold change between sample groups of at least 1.5-fold, ANOVA p values of <0.05, minimum CV of <30 % and a minimum normalized abundance of 10. Where putative compound annotations were present after statistical filtering, these were inputted into IMPaLA^51^ for metabolic pathway analysis.

### Whole genome sequencing

Genomic DNA was isolated from sorted cells or tail biopsies by using the QIAmp Fast DNA Tissue kit (#51404, Qiagen) according to the manufacturer’s protocol. Sequencing libraries were generated using on-bead tagmentation chemistry with the Illumina® DNA Prep, (M) Tagmentation Kit (Illumina) according to the manufacturer’s protocol. Briefly, bead-linked transposomes were used to mediate the simultaneous fragmentation of gDNA (100–500 ng) and the addition of Illumina sequencing primers. Next, reduced-cycle PCR amplification was used to amplify sequencing ready DNA fragments and to add the indices and adapters. Finally, sequencing-ready fragments were washed and pooled prior to paired-end sequencing (150 + 150 cycles, plus indices) on an Illumina NovaSeq6000 instrument. Finally, the output data were demultiplexed and BCL-to-Fastq conversion performed using Illumina’s bcl2fastq software, version 2.20.0.422.

Unmapped paired-reads of 151 bp were checked using a quality control pipeline including FastQC v0.11.3 and FastQ Screen v0.14.0. The reads were trimmed to remove any adapter sequence or poor-quality reads using Trimmomatic v0.39; reads were truncated at a sliding 4 bp window, starting 5’, with a mean quality <Q20, and removed if the final length was less than 35 bp. Only paired good quality reads were used in downstream analyses.

The filtered reads were mapped to the mouse strain FVB_NJ reference sequence (GCA_001624535.1) from the UCSC browser using BWA-MEM v0.7.17. The -M flag was used to flag secondary reads. Read group information was also added to each read. The mapped reads were further processed using samtools v1.17 to identify properly paired reads, fixmates, and sort the reads by coordinates. Picard Tools v2.27.5 MarkDuplicates was used to flag duplicated reads. InDels were detected in each sample using Manta v1.6.0. Strelka2 v2.9.10 was used to identify SNVs using Indels from Manta as input (via --indelCandidates).

SigProfilerMatrixGenerator v1.2.14 and SigProfilerExtractor v1.1.21 were used to generate reports on the context for SNV and InDel variants, and to match mutation signatures to COSMIC v3.3 GRCh37 profiles. The FVB_NJ mouse background was set up as custom genome in SigProfiler. The downloaded reference sequence was split into separate chromosomes, only retaining chr1-19, X and M. Transcript annotation was downloaded from Ensembl BioMart for Mus_musculus_FVB_NJ version 109.

### Cell line studies

Mouse mammary epithelial cell line, EpH4, was purchased from Sigma (#SCC284) and cultured in Dulbecco’s modified eagle medium (DMEM) (Sigma Aldrich, D6429), supplemented with 10% Fetal Bovine Serum (FBS) (Biosera, FB-1001/500) and 1% L-Glutamine (Sigma Aldrich, G7513). A transgenic EpH4 cell line with doxycycline-inducible HER2 overexpression was generated by lentiviral transduction using a lentiviral vector described in^52^. HER2 overexpression was induced in these transgenic cells with the addition of doxycycline (#D9891, Sigma) in their culture medium, which had been refreshed every 2 days of culture.

For immunoblot confirmation of transgenic HER2 overexpression, protein lysates were prepared using 1X RIPA buffer containing protease (#539131, Merck Millipore) and phosphatase (#524629, Merck Millipore) inhibitors. Western blotting was performed using protocols described earlier^18^ by using anti-HER2 (#2242, Cell Signaling) and anti-beta actin (ab8224, Abcam) primary antibodies. For visualizing ROS in these cells, CellROX Deep Red reagent (Invitrogen) was used according to the manufacturer’s protocols.

### Statistical analyses

Statistical tests appropriate for the analyzed datasets were selected based on population distribution, sample centrality/variability and data type to meet assumptions of tests and performed using GraphPad Prism software or other more specialized software mentioned in the corresponding methods sections above. P values <0.05 or Q values <0.05 (depending on the data type) were considered statistically significant. Data were expressed as mean ± standard deviation (SD) or mean ± standard error of the mean (SEM). Statistical approaches used for each individual data type are described in the corresponding methods section or in the corresponding figure legends.

### Data Availability

The data generated in this study are available in the publicly available data repositories with an embargo till the date of publication:

The mass spectrometry proteomics data have been deposited to the ProteomeXchange Consortium via the PRIDE partner repository with the dataset identifier PXD043315

The mass spectrometry metabolomics data have been deposited to the MetabolomicsWorkbench repository with Project ID: PR001714 at http://dx.doi.org/10.21228/M8542K

The whole genome sequencing data have been deposited to the ArrayExpress repository with the Accession ID: E-MTAB-13138

