## Supplementary Figures for "HER2 overexpression initiates breast tumorigenesis non-cell-autonomously by inducing oxidative stress in the tissue microenvironment"

**Figures S1 to S8**

### SUPPLEMENTARY FIGURE 1

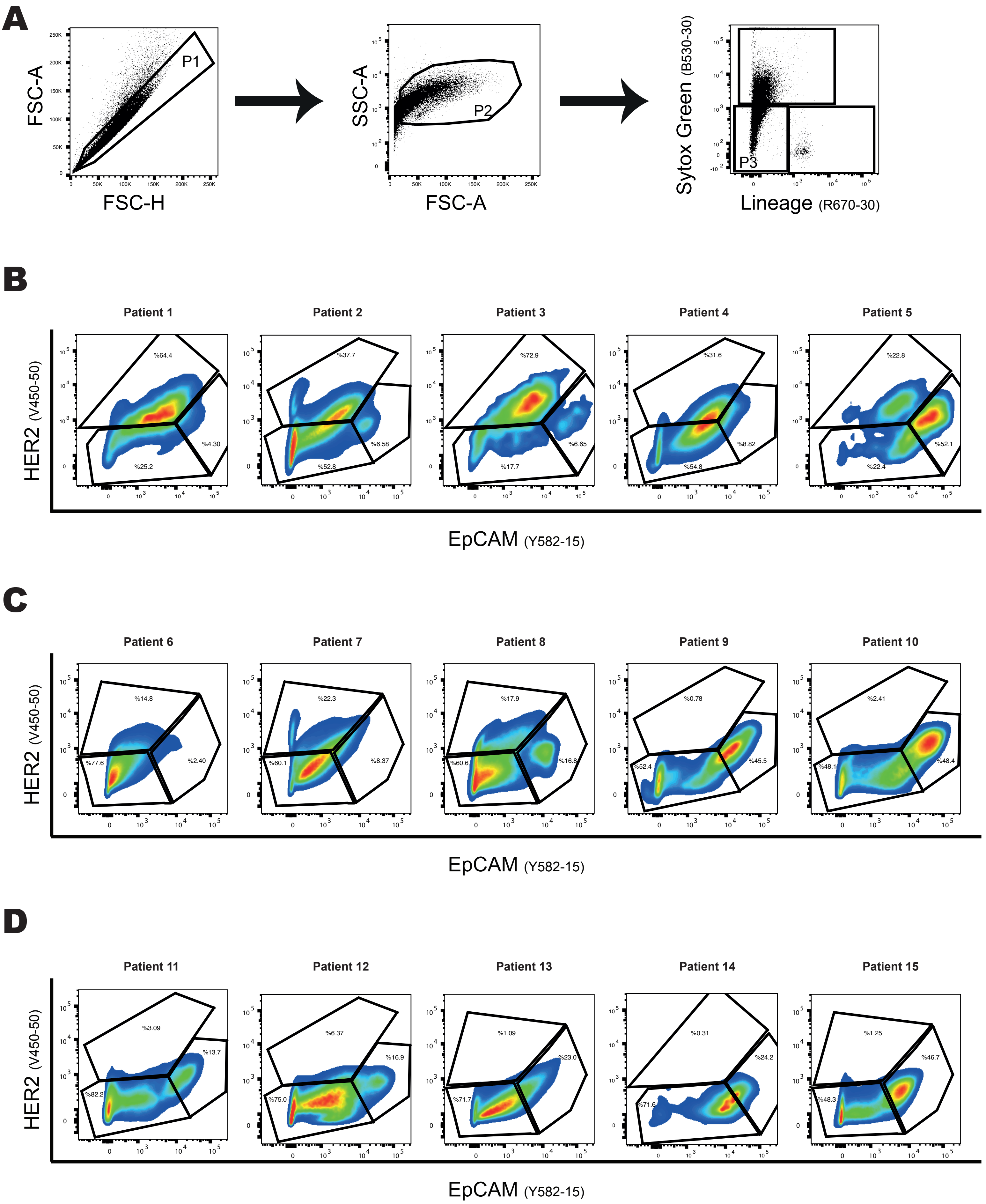

**Supplementary Figure 1.** Flow cytometry analysis of HER2 versus EpCAM expression in clinical samples of DCIS reveals the inter-patient heterogeneity

**A)** Representative dot plots demonstrate the gating strategy used for identifying DCIS cells. P1 gate identifies single cells based on forward-scatter values; P2 gate excludes the cell debris based on side- versus forward-scatter; P3 gate selects the alive (i.e negative for Sytox Green) and Lin$^{neg}$ (i.e not expressing CD45, CD31, or CD235A) cells, which are then analyzed for their HER2 versus EpCAM expression (**B-D**)

**B-D)** Pseudo-colored dot plots of 15 patient samples identifies three putative subgroups of DCIS based on the abundance of their HER2$^{pos}$ cells: (**B**) HER2-enriched group containing 40% or more HER2$^{pos}$ cells; (**C**) HER2-low group containing 15% to 30% HER2$^{pos}$ cells; and (D) HER2-negative group containing 10% or less HER2$^{pos}$ cells. In all groups, samples are ordered from left-to-right in descending order of the size of their EpCAM$^{hi}$ subpopulations. Of note, the plots in the middle of each row in **B-D** were shown as representative plots in Figure 1A.

### SUPPLEMENTARY FIGURE 2

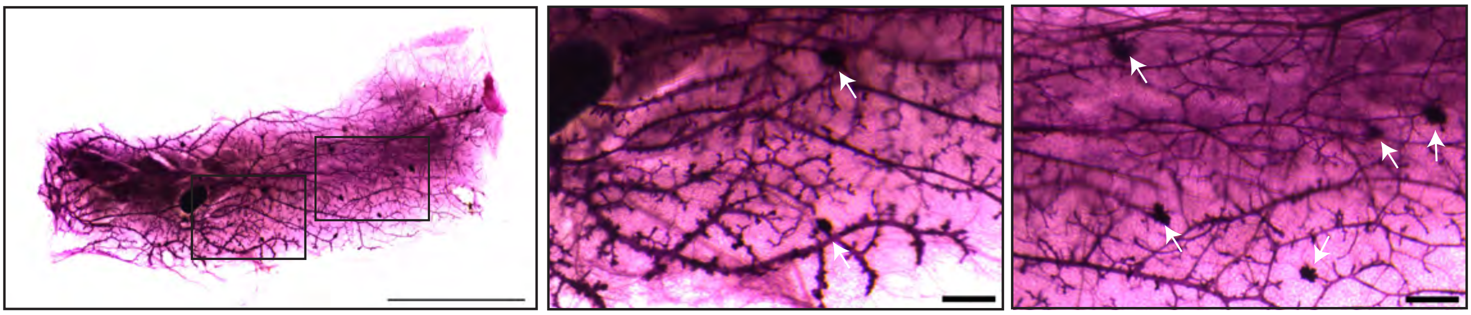

**Supplementary Figure 2.** Ductal lineage–dependency of early-stage mammary tumorigenesis does not occur when the onset of HER2 overexpression is at an adult age

Whole-mount staining results of No:4 inguinal glands of MMTV-rTTa;TetON-HER2 double-transgenic females that received doxycycline-containing food for 4 months starting from 15 weeks old age. Images shown are representative of n=3 mice. Multi-nodal early-stage tumors, which were randomly distributed within the mammary ductal tree, are shown with arrows. Transparent rectangles in the leftmost image point to the locations of the gland shown by the higher-magnification images. Scale bars represent 1 cm for the leftmost image and 1 mm for the other high-magnification images.

### SUPPLEMENTARY FIGURE 3

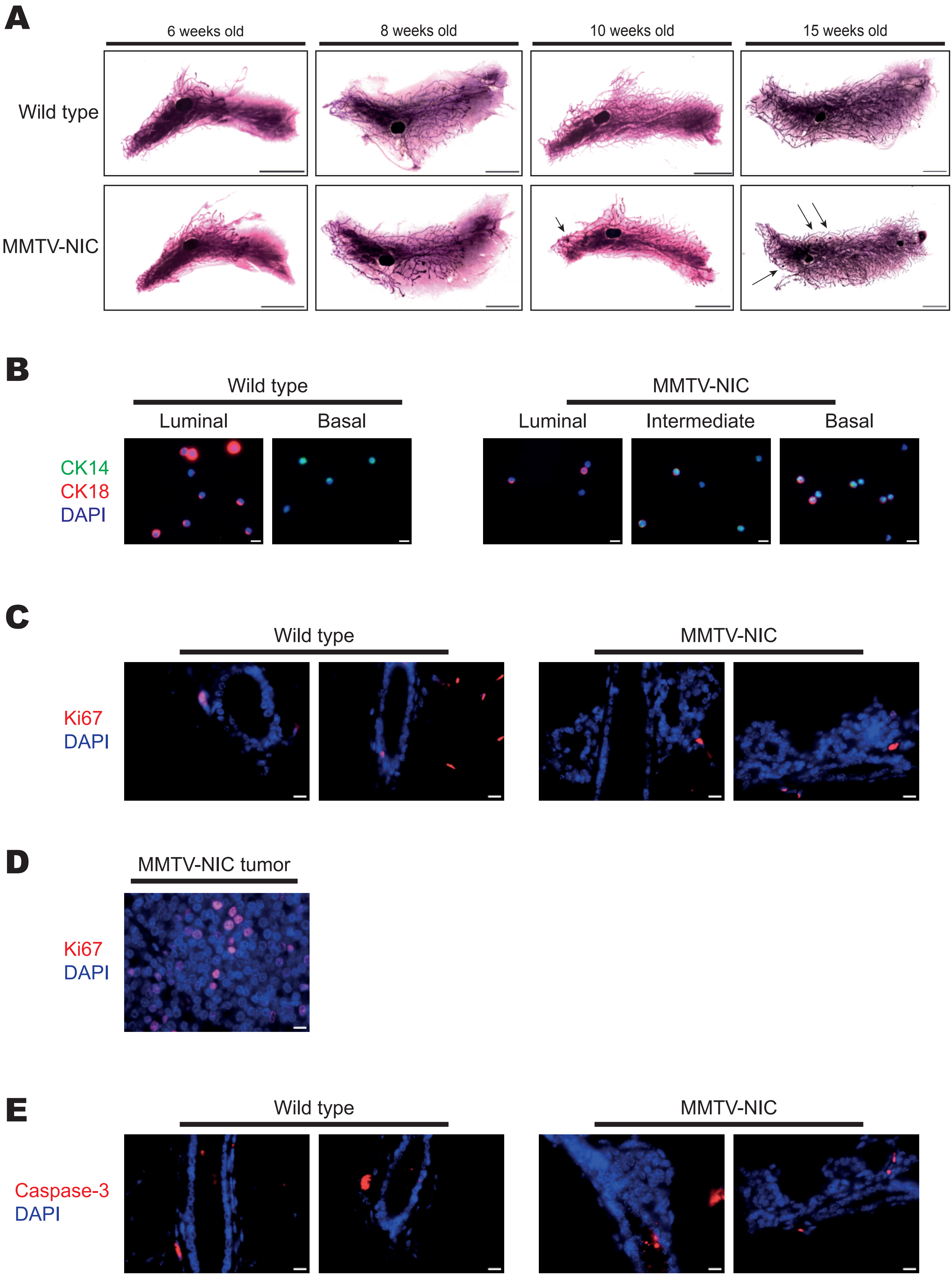

**Supplementary Figure 3.** Mammary glands of 8-week-old MMTV-NIC mice are free of tumors but contain ectatic ducts that are composed of mammary epithelial cells without increased levels of proliferation or apoptosis markers

**A)** Representative images of whole-mount staining results of No:4 inguinal glands of MMTV-NIC females at 6, 8, 10, or 15 weeks of age and their wildtype littermates. Arrows point to the early stage tumors in mammary glands of 10- and 15-week-old MMTV-NIC mice. Images shown are representative of at least 3 mice per genotype and age group. Scale bars represent 1 cm.

**B)** Representative images of mammary epithelial cells that were cytospun and co-stained with CK14, CK18, and DAPI. Images shown are representative of cells isolated from n=3 mice per genotype. Quantification results of this experiment are shown in Figure 4D.

**C)** Immunofluorescence staining of microtome sections of No:4 glands from 8-week-old MMTV-NIC and wildtype littermate mice for Ki67 shows no obvious increase in proliferation within ectatic ducts. Images shown are representative of 3 pairs of mice analyzed. Scale bars represent 10 um.

**D)** Immunofluorescence staining of microtome sections of a MMTV-NIC tumor for Ki67 shows increased numbers of proliferating cells within tumors. Scale bars represent 10 um.

**E)** Immunofluorescence staining of microtome sections of No:4 glands from 8-week-old MMTV-NIC and wildtype littermate mice for cleaved Caspase-3 shows no obvious increase in apoptotic cell numbers within ectatic ducts. Images shown are representative of 3 pairs of mice analyzed. Scale bars represent 10 um.

### SUPPLEMENTARY FIGURE 4

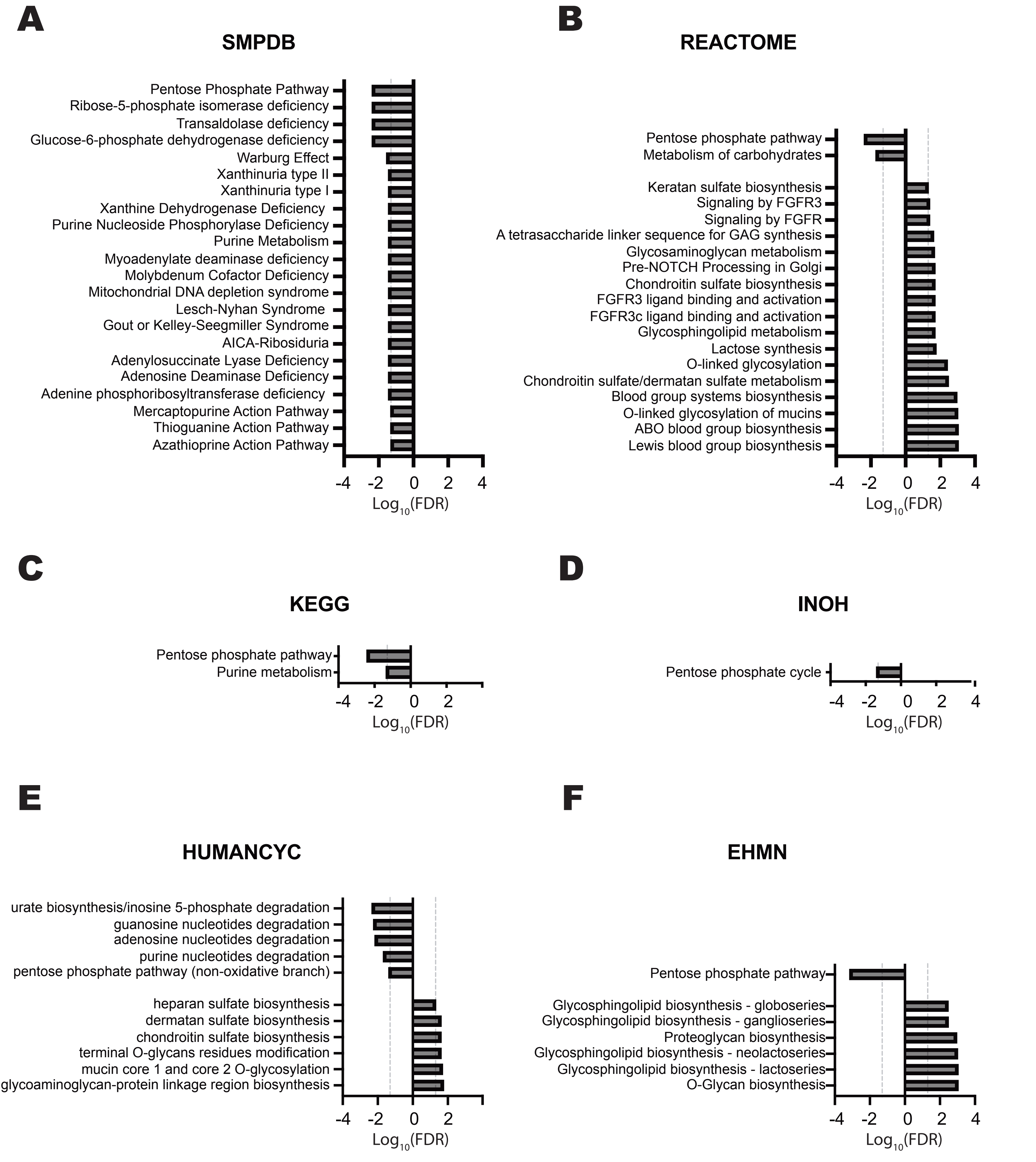


**Supplementary Figure 4.** MMTV-Neu mammary ducts have altered metabolic activities
**A-F)** Metabolomics dataset for intact mammary ducts of 8-week-old MMTV-NIC mice compared with their wildtype littermates was compared against 6 databases for pathway enrichment analysis. The dysregulated metabolic activities indicated by SMPDB (**A**), Reactome (**B**), KEGG (**C**), INOH (**D**), HUMANCYC (**E**), and EHMN (**F**) databases are shown as individual graphs. Vertical dashed lines represent the border of q=0.05. Metabolic activities that are downregulated or upregulated are represented as bars towards left from midline or towards right from midline, respectively.

### SUPPLEMENTARY FIGURE 5

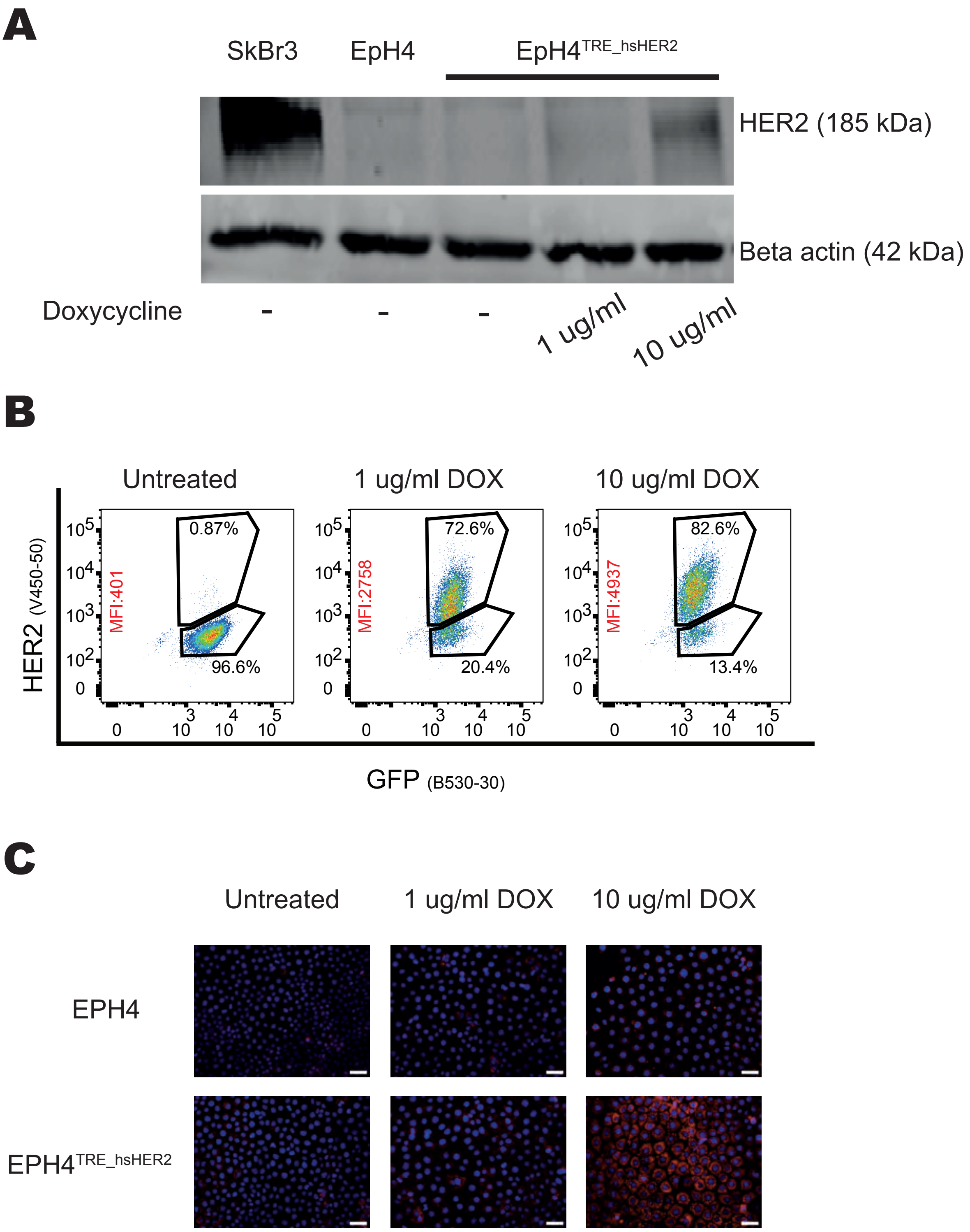


**Supplementary Figure 5.** Overexpression of HER2 in EpH4 cells results in increased ROS levels

**A)** Immunoblot analyses of human wildtype HER2 overexpression in transgenic EPH4 cells with doxycycline-inducible HER2 overexpression (i.e. EpH4$^{TRE_hsHER2}$). The HER2+ breast cancer cell line, SkBr3 was used as a positive control. The transgene expression was induced by treatment with doxycycline at the indicated concentrations. Beta actin was used as the loading control.

**B)** Representative dot plots of flow cytometry analysis of EpH4$^{TRE_hsHER2}$ cells for the surface expression of HER2 upon treatment with doxycycline to induce transgene expression at the indicated concentrations. The mean fluorescence intensity (MFI) for HER2 expression is shown on the dot plots (text in red). Of note, in these cells the GFP expression marks the transduced cells but is not dependent on the doxycycline treatment. Shown plots are representative of three independent experiments.

**C)** CellROX Deep Red staining of parental EpH4 and EpH4$^{TRE_hsHER2}$ cells upon treatment with the indicated concentrations of doxycycline for 7 days in culture demonstrates increased levels of ROS accumulation (in red) upon HER2 overexpression. Cells were counterstained with DAPI (in blue). Scale bars represent 50 um.

### SUPPLEMENTARY FIGURE 6

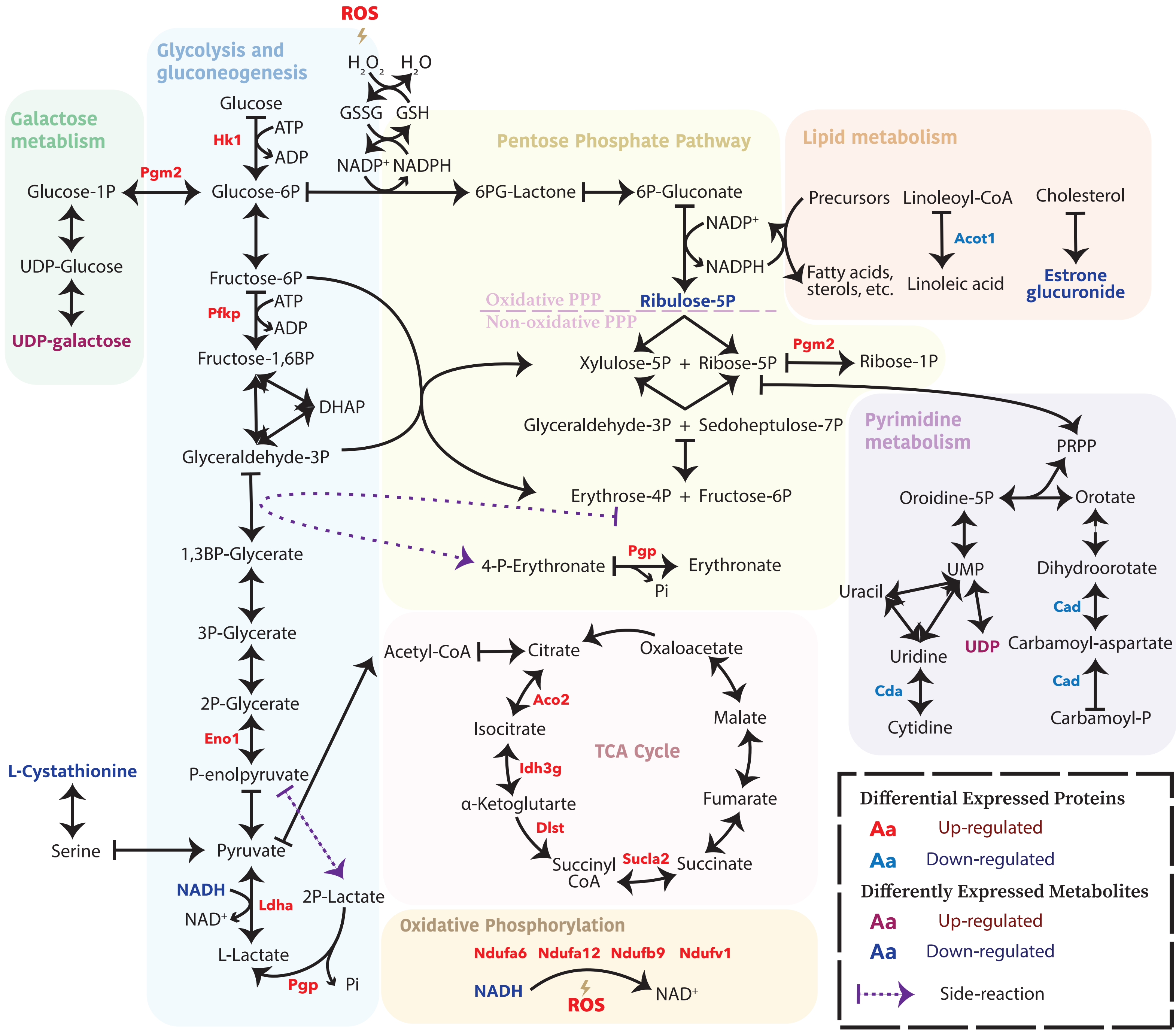


**Supplementary Figure 6.** Integrated network analysis of the proteome and metabolome datasets for the MMTV-NIC mammary ducts illustrates the inter-connections between altered metabolic pathways

The key metabolites of central carbon metabolism are shown for glycolysis, pentose-phosphate pathway, and TCA cycle along with simplified versions for metabolites in oxidative phosphorylation, pyrimidine metabolism and lipid metabolism that are relevant to the findings of this study. Metabolites with significant ($q<0.05$) alterations in MMTV-NIC ducts compared to wildtype ducts are marked. Proteins/enzymes involved in these metabolic pathways and demonstrated significant ($p<0.05$) alterations in MMTV-NIC ducts are named near the arrows representing the corresponding enzymatic activity.

### SUPPLEMENTARY FIGURE 7

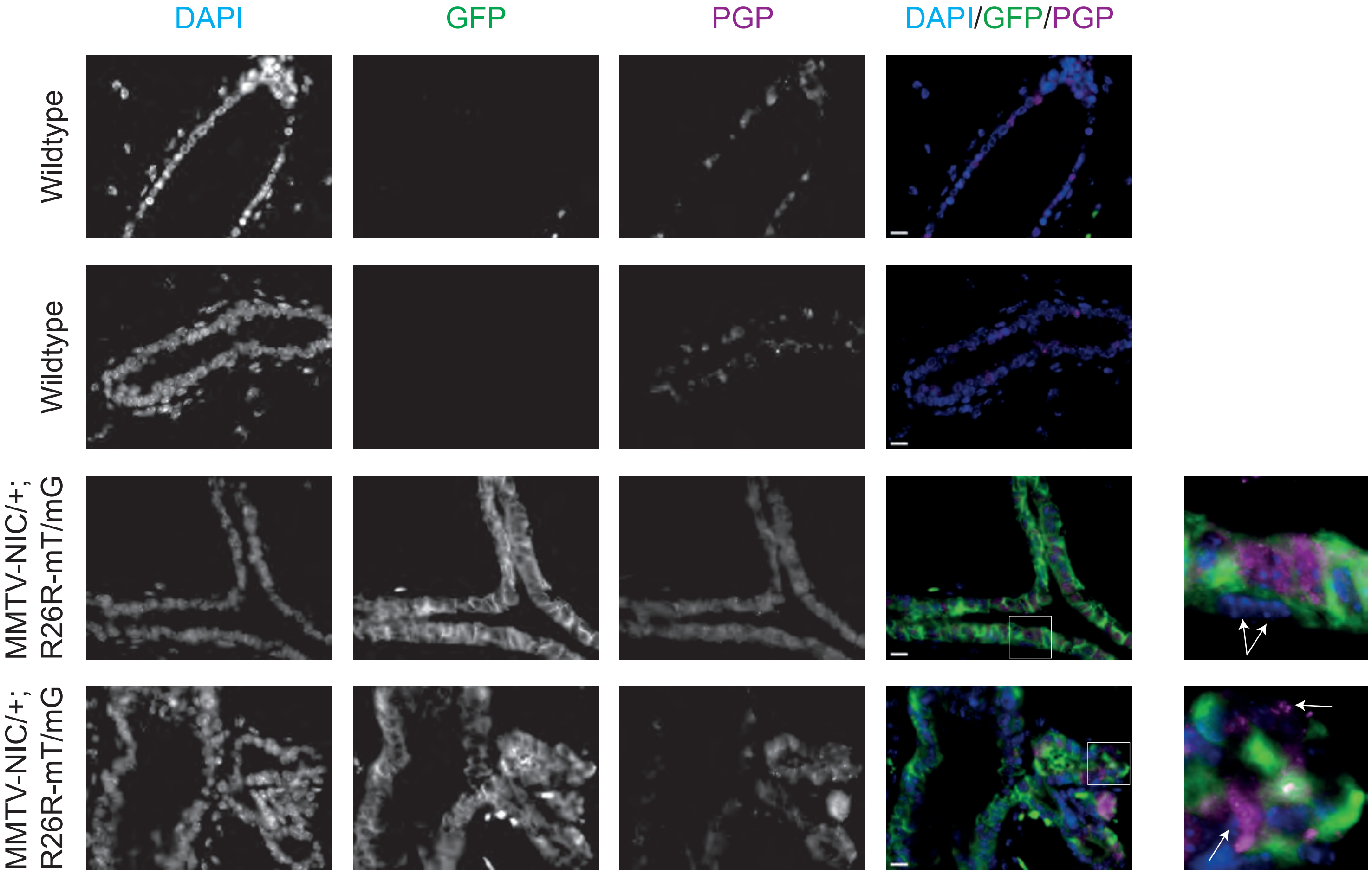


**Supplementary Figure 7.** PGP is expressed in Neu$^{neg}$ lineage cells only in the ectatic ducts of the MMTV-Neu glands

Co-immunofluorescence staining of microtome sections of No:4 glands from 8-week-old MMTV-NIC;R26R-mT/mG and wildtype littermate mice for the expression of GFP -the marker of the Neu$^{pos}$ lineage- versus PGP. In bilayered ductal structures there is a low level of PGP expression only in Neu$^{pos}$ lineage cells; whereas in ectatic ducts a high level of PGP expression was observed in both Neu$^{pos}$ and Neu$^{neg}$ lineage cells. Arrows in the high-magnification images point to the Neu$^{neg}$ lineage cells. Images shown are representative of 3 pairs of mice analyzed. Scale bars represent 10 um.

### SUPPLEMENTARY FIGURE 8

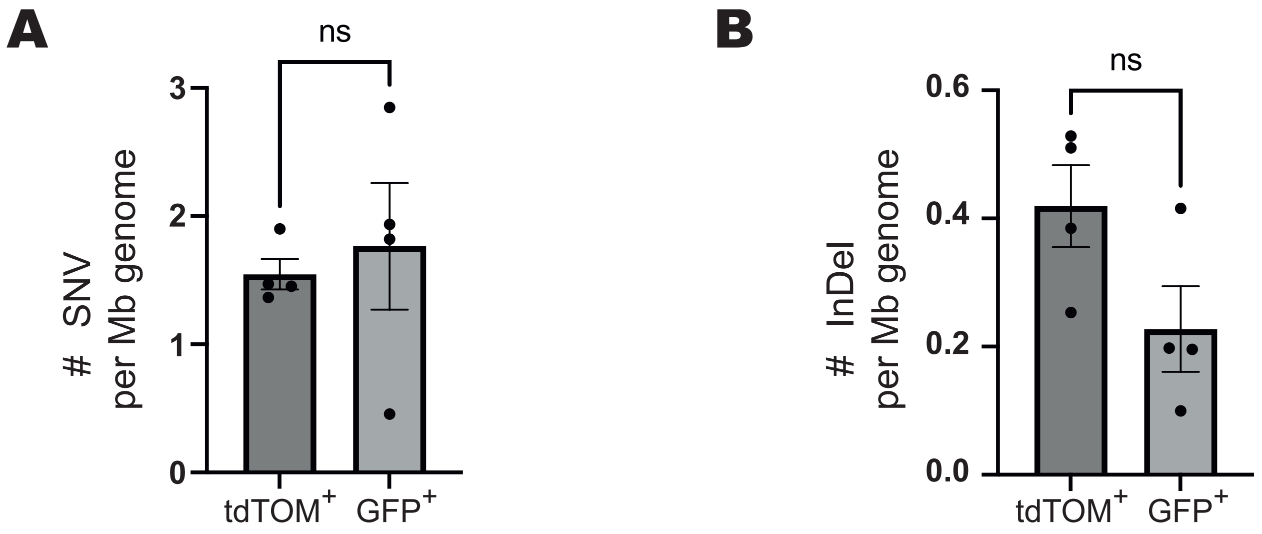


**Supplementary Figure 8**. Tumor cells within the $Neu^{neg}$ and $Neu^{pos}$ lineages have similar levels of mutation burden

Average numbers of SNVs (**A**) or InDels (**B**) per Mb of genome were similar for the $tdTOM^{+}$ $Neu^{neg}$ versus $GFP^{+}$ $Neu^{pos}$ lineage tumor cells. The graphs represent mean ± SEM of 4 tumors analyzed with individual values shown. Differences between sample groups were not significant according to two-tailed paired t-test.

.
